# A preliminary study of resting brain metabolism in treatment-resistant depression before and after treatment with olanzapine-fluoxetine combination

**DOI:** 10.1101/624288

**Authors:** José V. Pardo, Sohail A. Sheikh, Graeme Schwindt, Joel T. Lee, David E. Adson, Barry Rittberg, Faruk S. Abuzzahab

## Abstract

Treatment-resistant depression (TRD) occurs in many patients and causes high morbidity and mortality. Because TRD subjects are particularly difficult to study especially longitudinally, biological data remain very limited. In a preliminary study to judge feasibility and power, 25 TRD patients were referred from specialty psychiatric practices. All were severely and chronically depressed and mostly had comorbid psychiatric disorders as is typical in TRD. Nine patients were able to complete all required components of the protocol that included diagnostic interview; rating scales; clinical magnetic resonance imaging; medication washout; treatment with maximally tolerated olanzapine-fluoxetine combination for 8 weeks; and pre- and post-treatment fluorodeoxyglucose positron emission tomography. This drug combination is an accepted standard of treatment for TRD. Dropouts arose from worsening depression, insomnia, and anxiety. One patient remitted; three responded. A priori regions of interest included the amygdala and subgenual cingulate cortex (sgACC; BA25). Responders showed decreased metabolism with treatment in the right amygdala that correlated with clinical response; no significant changes in BA25; better response to treatment the higher the baseline BA25 metabolism; and decreased right ventromedial prefrontal metabolism (VMPFC; broader than BA25) with treatment which did not correlate with depression scores. The baseline metabolism of all individuals showed heterogeneous patterns when compared to a normative metabolic database. Although preliminary given the sample size, this study highlights several issues important for future work: marked dropout rate in this study design; need for large sample size for adequate power; baseline metabolic heterogeneity of TRD requiring careful subject characterization for future studies of interventions; relationship of amygdala activity decreases with response; and the relationship between baseline sgACC and VMPFC activity with response. Successful treatment of TRD with olanzapine-fluoxetine combination shows changes in cerebral metabolism similar to those seen in treatment-responsive major depression.

## Introduction

Treatment-resistant depression (TRD) is often operationally defined as a major depressive episode that fails to remit after treatment with at least two antidepressants of different classes at therapeutic doses for an adequate treatment period [1-8]. More extensive staging criteria for TRD have been proposed as well [1]. TRD must be distinguished from inadequately treated depression resulting from numerous factors such as patient non-compliance, intolerance to side effects, misdiagnosis (e.g., thyroid disease), low dosage, etc. [2]. It is unclear if TRD is a particularly malignant form of depression with its own pathophysiology, or if treatment-related changes in brain metabolism are different than those found in treatment-responsive depression [3]. Patients with TRD are at increased risk of relapse [4]. Also, most TRD patients have numerous psychiatric comorbidities inherently raising potential confounds with diagnosis [5-8].

The STAR*D trial documents TRD is a frequent occurrence and a serious problem in psychiatry afflicting about 30% of patients [4]. In terms of disease burden, it is second only to back pain in terms of life-years of disability [9]. The high significance of TRD has prompted an aggressive search for novel, more effective treatments including new classes of antidepressants (e.g., glutamatergic receptor antagonists, neuroactive steroids), add-on treatments (drug combinations, boosters such as lithium or liothyronine, atypical neuroleptics, mood stabilizers), and devices for neuromodulation (e.g., transcranial magnetic stimulation, vagus nerve stimulation, deep brain stimulation, direct current stimulation).

Despite the clinical importance of TRD, few studies have examined its biology and treatment. Several challenges have hindered such research. The recruitment of TRD patients is difficult. These patients are likely heterogenous in pathology, have numerous comorbidities, and are quite ill with significant risk for suicide. Their cross-sectional physiology may be confounded by the many previous treatment trials. At best, the medley of ineffective medications needs wash out, but this may lead to potential withdrawal symptoms or symptom worsening. Encouragement to undergo yet another treatment after so many failed trials becomes paramount. These patients need frequent follow-up and clinician availability. Despite these impediments, some work has been done using F^18^-fluorodeoxyglucose positron emission tomography (FDG PET) in those with TRD [10-18]. However, not only does the biology of TRD, if homogeneous, remain unclear, but also no biomarkers predicting treatment resistance have reached clinical utility for this group of patients.

Past neuroimaging studies of treatment-responsive depression have highlighted the amygdala and subgenual anterior cingulate cortex (sgACC) as key nodes of depression-related circuity showing reduction in activity with successful treatment (see reviews [19-21]), although there are exceptions (e.g., no sg ACC change [22]; no amygdala change [23]). Studies have suggested several features may characterize TRD such as sgACC hyperactivity, amygdala hyperactivity, prefrontal/thalamic dysconnectivity, habenular connectivity, prefrontal hypoactivity, and hippocampal subfield volumes [10, 13, 14, 24-26]; however, consensus is yet to be achieved.

To our knowledge, there are no prior studies of brain metabolism in TRD patients with drug washout to establish a baseline examined both before and after a full trial of a combination antidepressant and atypical neuroleptic. Yet, the combination of antidepressant and atypical neuroleptic is being used increasingly throughout the world to treat TRD. The present study is of necessity preliminary, as no prior data existed with which to power the sample size or even to determine its feasibility.

With a focus on the amygdala and sgACC based on prior literature, this cohort observational study sought to test feasibility and to characterize regional brain metabolism in TRD before and after adequate treatment with a combination of an atypical neuroleptic (olanzapine) and fluoxetine (O/F), a selective serotonin reuptake inhibitor (SSRI) antidepressant. Use of these drugs in combination will be referred to hereafter as O/F.

This drug combination was the first drug approved by the USA Food and Drug Administration in 2009 specifically for the indication of TRD [27]. There have been several clinical trials, reviews, and meta-analyses examining the efficacy of O/F for TRD; that discussion is beyond the scope of this study [28-35]. Originally, use of O/F was based on preclinical work. Neurobiological changes associated with O/F dosing in rats include increased prefrontal monoamine levels [36, 37] and suppression of limbic immediate-early gene expression [38] relative to olanzapine or fluoxetine alone. No differential effects on limbic neurogenesis were found using O/F, whereas the individual drugs are associated with neurogenesis [39]. Whereas higher doses of O/F increased levels of neurotrophin-3 selectively in rat prefrontal cortex, low O/F doses or higher doses of olanzapine or fluoxetine administered individually did not [40]. These animal findings suggest some unique effects of O/F therapy not accounted for by the actions of the individual drugs.

The aim of the present work was not to assess efficacy or health outcomes in a clinical trial, as this has been reported previously (see above). Rather, the project’s purpose was to use imaging to bear on the question of mechanisms. Whether such combination treatments for TRD follow similar metabolic effects to other antidepressants in treatment-responsive depression is unknown. In addition, the individual TRD patient’s deviation from a normative database indicate for the first time the potential changes in regional brain metabolism in an individual, unmedicated, TRD patient. Such data could address preliminarily whether sgACC hypermetabolism or other biomarker is characteristic of TRD, a key issue in patient selection for future treatment trials of TRD.

## Materials and Methods

### Participants

Twenty-five participants with severe TRD were enrolled. They had many psychotherapy and medication trials, some even failing convulsive therapy. They all had a longstanding chronic illness lasting many years. Most had comorbid psychiatric disorders. They were recruited and enrolled through referral from physicians’ outpatient clinics known to specialize in the treatment of TRD (co-authors: DA, BR, FSA). The principal inclusion criterion was severe, refractory major unipolar depressive disorder (Scheduled Clinical Interview for Diagnostic and Statistical Manual-IV; SCID [41]) as the primary diagnosis. Exclusion criteria included a lifetime history of cognitive impairment, psychosis, bipolar disorder, drug dependence, pregnancy, as well as any clinically significant findings on magnetic resonance imaging. All subjects provided written informed consent as approved by the VA Institutional Review Board (IRB) and the Radioactive Drug Research Committee (RDRC) approved by the FDA.

### Treatment

Patients’ polypharmacy was tapered with washout for two weeks before the baseline measurement of glucose uptake using FDG PET as described previously [42]. They were then titrated to the maximal tolerated dose of fluoxetine (≤ 60 mg) and olanzapine (≤ 30 mg). The use of the combination of an SSRI such as fluoxetine and an atypical neuroleptic such as olanzapine is a standard of care in the management of patients with TRD. The maximal tolerated dose was held constant for eight weeks except for rescue medication of low dose benzodiazepine or short acting hypnotics for severe anxiety or insomnia. Given the seriousness of the illness, risk for suicide, and focus on mechanisms rather than efficacy, the study had no placebo control medication. Patients were seen either weekly or every two weeks (depending on stability) during wash out and treatment. Compliance was checked by counting of pills every 1-2 weeks. Side effects were evaluated with open-ended questions without checklists typical of clinical trials. After treatment with O/F for eight weeks and PET imaging, they were returned to their referring psychiatrist for continued assessment and follow-up.

### Clinical Assessments

All subjects were assessed by their referring physician as having TRD as their primary diagnosis. Medical records were reviewed to ensure all subjects had at least two trials of antidepressants from different classes with adequate doses and duration of treatment (at least stage III TRD [1]). All subjects underwent structured diagnostic interviews using the SCID-1, Clinician Version [43]. The primary outcome measure was the change in Montgomery-Asberg Depression Rating Scale (MADRS [44]) score after eight weeks of treatment. A clinical response was defined as a greater than or equal to 50% drop in depression score, while a remission was defined as a MADRS of less than or equal to 8. Anxiety was scored with the Hamilton Anxiety Scale (HAMA [45]). Additional testing not directly pertinent to the present study included Clinical Global Impression Scale (CGI [46]), Mini-Mental Status Exam (MMSE [47]), Shipley Institute of Living Scale [48], Edinburgh Handedness Inventory [49], Profile of Mood States (POMS) [50], and Positive Affect Negative Affect Scale (PANAS [51]).

### Positron emission tomography

Patients were scanned after washout (pre-treatment or baseline) and after completion of the eight weeks of treatment at maximal tolerated dose (post-treatment). Participants fasted for at least six hours before imaging; blood glucose was checked immediately before scanning. The relative regional glucose uptake was measured by injecting intravenously a bolus of ^18^F-FDG in saline at a dose of 185 MBq (5 mCi)/70 kg. They rested for 50 minutes during tracer uptake with eyes closed and ears open in a dimly lit, quiet room while being monitoring for wakefulness. The scanner was a Siemens (Knoxville, TN, USA) ECAT EXACT 47 operated in 2-D mode with septae extended. After measured attenuation, counts were collected during an emission scan lasting 20 minutes. Data were corrected for scatter, decay, randoms, and electronic deadtime. Images were reconstructed using filtered backprojection to a resolution of approximately 12 mm full width at half maximum.

### Image analysis

Image analysis followed routine procedures including normalization for whole-brain activity, intersubject stereotactic averaging, subtraction of pre- from post-treatment activity, statistical parametric mapping, and threshold Z = 3.3 as previously described [52]. Images were warped nonlinearly into stereotactic space [53] with regression for age using Neurostat software [54]. Parametric maps were overlaid on a template MRI blurred to a similar resolution as the PET scan. In-house software (iiV, http://james.psych.umn.edu/iiV/) was used to display results on a standard MRI template [55]. For an exploratory look at individual TRD patient’s whole-brain voxelwise differences from our normative database (N = 30). For the purposes of display, the individually warped *t* images used an uncorrected magnitude threshold of p < ~0.05.

### Regions of interest

Two ROIs were examined based upon existing, extensive literature (see Fig 1): amygdala and sgACC. The amygdala ROI consisted of a sphere of 13 mm diameter center on each amygdala in the atlas of Talairach and Tournoux (1988, Fig 1) at coordinates (±23, −4, −16). Mean counts were collected for each ROI from the pre- and post-treatment scans for each patient. The percent change in MADRS score was regressed linearly against the mean amygdala activity. Paired t-tests were used to compare mean amygdala counts before and after treatment for the group and for responders vs. non-responders separately. A threshold of p < 0.05 was used for the ROI analysis.

**Figure 1.**
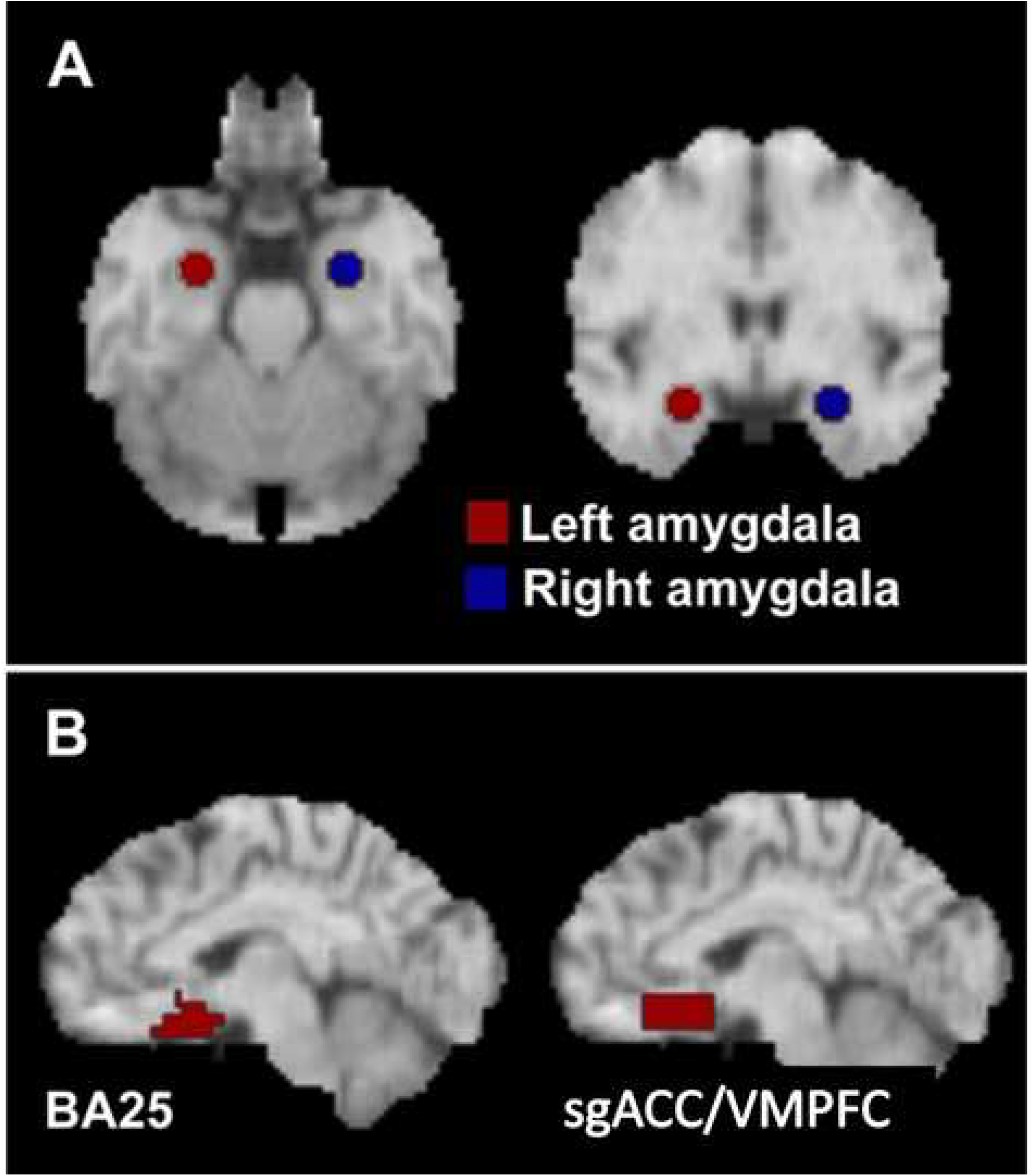
Amygdala (A) and sgACC/VMPFC (B) regions of interest.

Based upon an *a priori* focus on the sgACC, we defined two additional subregions within sgACC for this analysis. Previous studies point to changes in subgenual metabolism that are not restricted to the sgACC of Brodmann area 25 (BA25) but extend along the ventromedial cingulate cortex [10, 18, 56, 57]. So, a region including only those voxels labeled as BA25 by the atlas of Talairach and Tournoux [53] was created using the Talairach Daemon (http://ric.uthscsa.edu/projects/talairachdaemon.html) [58]. A second, less specific region was drawn as a cuboid on each hemisphere with the following extents measured from the anterior commissure: x, ±(1-12 mm); y, 10-25 mm; z, −5 to −16) mm [53]. This approximated the region of hypometabolism associated with antidepressant treatment reported previously by our laboratory [18]. The region encompasses BA25, as well as portions of BA32 and BA33 and will be referred to as subgenual/VMPFC (Fig 1B).

Mean counts from each of these subgenual regions were compared pre- and post-treatment for responders and non-responders using paired *t*-tests. Percent-change in mean ROI metabolism was linearly regressed against percent-change in MADRS across all subjects. Pre-treatment mean metabolism was also regressed against percent change in MADRS.

One additional unplanned ROI was included post hoc because of its proximity to the amygdala, involvement in depression, and similar response to treatment in published work [22]. A 13 mm diameter spherical ROI was centered on the hippocampus at the following Talairach coordinates: x, ±27 mm, y, −23 mm; z, −9 mm or MNI (±27, −22, −0) http://sprout022.sprout.yale.edu/mni2tal/mni2tal.html.

A repeated measures ANOVA was performed on the amygdala, VMPFC, and head of the hippocampus ROIs as defined above. Extensive prior literature highlights these regions as important in affective illness. The BA25 region was not included as no change in activity occurred with treatment (see below). The dependent measure was glucose uptake (PET counts). Repeated measures included TIME (Pre-treatment, Post-treatment), ROI (amygdala, hippocampus, and VMPFC), and SIDE (right hemisphere, left hemisphere). Additionally, the correlation matrix for glucose uptake across amygdala, VMPFC, and hippocampus was calculated (S2 Table).

## Results

### Clinical response

Drug wash out led frequently to increased insomnia, anxiety, and depression which were the leading reasons for discontinuation by participants (N = 16). Side effects from O/F observed reflected those commonly seen with this drug combination [28]. There were no suicide attempts during the study. Nine participants completed the entire protocol (clinical MRI; drug washout; O/F treatment; Pre- and Post-treatment PET scans). Clinical data are shown in Table 1 which includes demographics, gender, family history of depression, illness onset, comorbidities (both current and lifetime), failed treatments, and response designation. The HAMA and MADRS scores as well as weights before and after treatment are displayed in S1 Table. The average tolerated dose was 39 mg (range 30-45 mg) of fluoxetine and 12 mg (range 10-12 mg) of olanzapine. Their weight increased significantly during treatment (S1 Table; S1 Figure); there was no interaction between time (pre vs post) and response (*p* > 0.2). The average baseline MADRS score was 31 (SD 5; range 24-38); no significant differences arose in baseline MADRS score between responders vs. non-responders (*p* > 0.3; no shown). Completers showed a decrease in MADRS score following eight weeks of treatment (Mean, 12; SD, 5; S1 Table). The baseline HAMA did not differ between responders vs. non-responders (*p* < 0.72). The HAMA declined significantly also after treatment (*t*(8) = 4.7; *p* < 0.002) with the significance driven by the non-responders (*p* < 0.006; responders, *p* < 0.13). Four patients responded; one of these achieved remission. Of note, responders tended to have fewer comorbidities and earlier onset than non-responders.

**Table 1.**
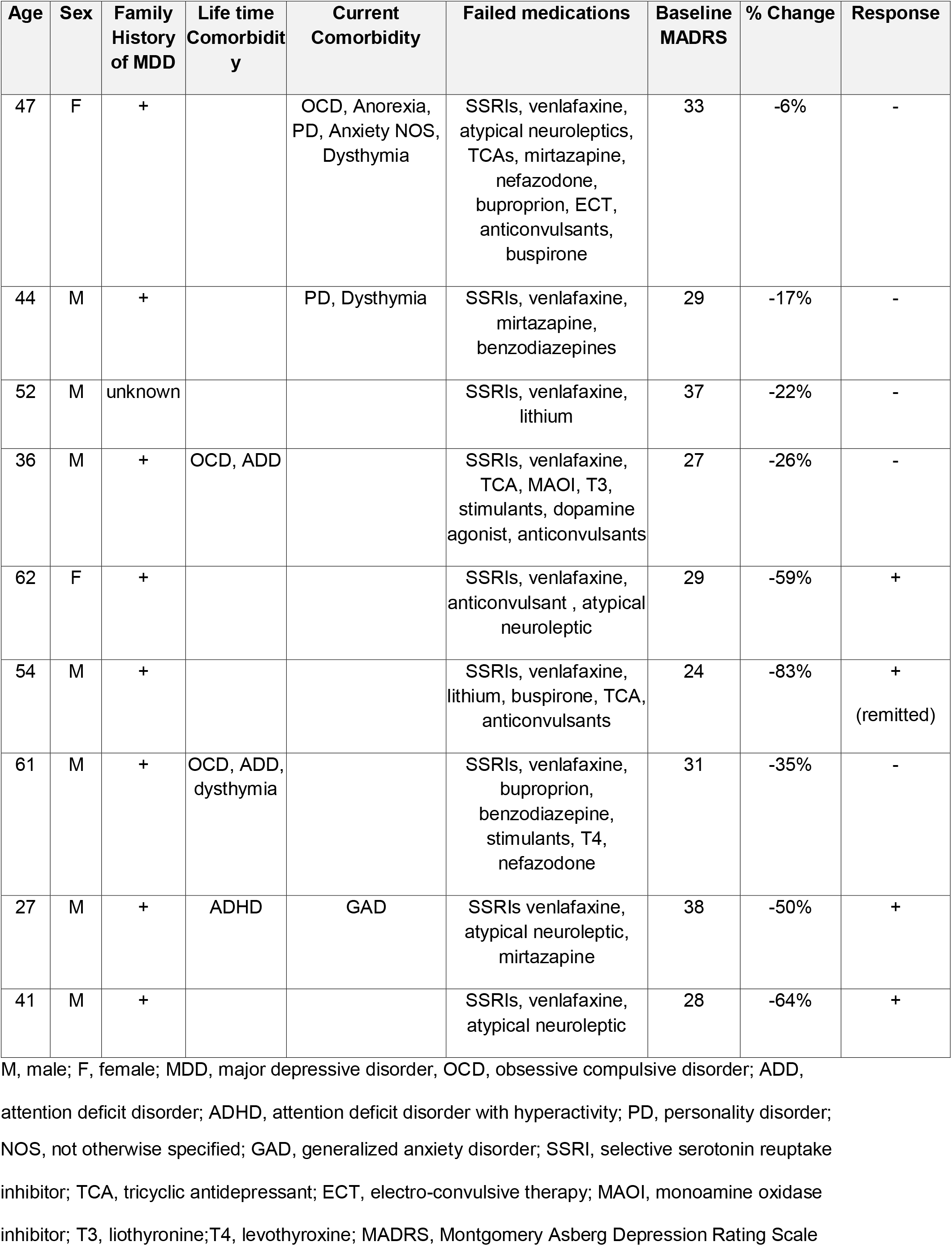
Patient demographics and outcomes.

A 2 x 2 Analysis-of-Variance with Treatment-response (responder vs. non-responder) as the between-subject factor and Time (post vs. pre-test) as the within-subject factor demonstrated a main effect of Time (*F*(1,7) =195.2, *p* < 0.001), confirming that MADRS scores decreased significantly across sessions. The interaction of Treatment-response X Time was also significant (*F*(1,7) = 43.9 *p* < 0.001). This interaction confirmed that responders showed significantly greater reductions in MADRS scores than non-responders.

### Whole-brain exploratory image analysis

Voxel-wise comparisons of patients’ post- and pre-treatment scans revealed significant changes (Table 2; S2-4 Figures). Patients were grouped as responders and non-responders (see above) to identify metabolic change associated with successful treatment.

**Table 2.** Significant changes in whole-brain voxelwise analysis.

| X<br>(mm) | Y<br>(mm) | Z<br>(mm) | Z-score | Region |
| --- | --- | --- | --- | --- |
| **All patients** |  |  |  |  |
| 12 | -35 | 54 | +3.4 | R paracentral lobule |
| 6 | -60 | 43 | +3.3 | R precuneus (BA7) |
| -12 | -60 | -20 | -4.6 | L Cerebellum |
| 17 | -94 | -4 | -3.7 | R Lingual Gyrus (BA18) |
| 28 | 14 | -20 | -3.6 | R Inferior Frontal Gyrus (BA47) |
| -15 | -33 | 7 | -3.6 | L Thalamus |
| -37 | -64 | -22 | -3.5 | L Cerebellum |
| 30 | -49 | -27 | -3.4 | R Cerebellum |
| -12 | -10 | 0 | -3.4 | L Thalamus |
| 10 | -1 | -4 | -3.3 | R globus pallidus |
| **Non-responders** |  |  |  |  |
| -8 | -60 | -16 | -5.2 | L Cerebellum |
| 8 | -49 | -29 | -4.6 | R Cerebellum |
| -17 | -53 | 29 | -4.4 | L Cingulate Gyrus (BA31) |
| -44 | 32 | 7 | -3.9 | L Inferior Frontal Gyrus (BA45) |
| -10 | -10 | 2 | -3.9 | L Thalamus |
| 17 | -91 | -2 | -3.7 | R Lingual Gyrus (BA17) |
| -51 | -67 | 14 | -3.7 | L Middle Temporal Gyrus (BA39) |
| -21 | -73 | -18 | -3.7 | L Cerebellum |
| 1 | -19 | -16 | -3.4 | R Midbrain |
| -42 | -4 | 38 | -3.4 | L Precentral Gyrus (BA6) |
| **Responders** |  |  |  |  |
| -48 | -13 | 2 | +3.5 | Left Superior Temporal Gyrus (BA22) |
| -53 | -24 | 18 | +3.4 | Left Post-Central Gyrus (BA40) |
| 6 | -49 | 36 | +3.3 | Right Precuneus (BA31) |
| 10 | -58 | 38 | +3.3 | Right Precuneus (BA7) |
| 19 | -1 | -7 | -4.4 | Right Dorsal Amygdala |

R, right; L, left; BA, Brodmann area

Responders showed increases in the left superior temporal and post-central gyri and right precuneus. Significantly reduced metabolism was confined to a peak in the right amygdaloid complex (see Fig 2). Of note, liberalizing the threshold to 0 < 0.05 (uncorrected) revealed bilateral amygdala deactivations. Non-responders showed no significant increases in metabolism following treatment. Areas of reduced metabolism after treatment were found in non-responders in the bilateral cerebellum, lingual gyrus, middle temporal gyrus, thalamus, midbrain, and dorsal cingulate gyrus (see Table 2). For all patients as a group, no significant changes occurred in limbic structures in the whole-brain image analysis.

**Figure 2.**
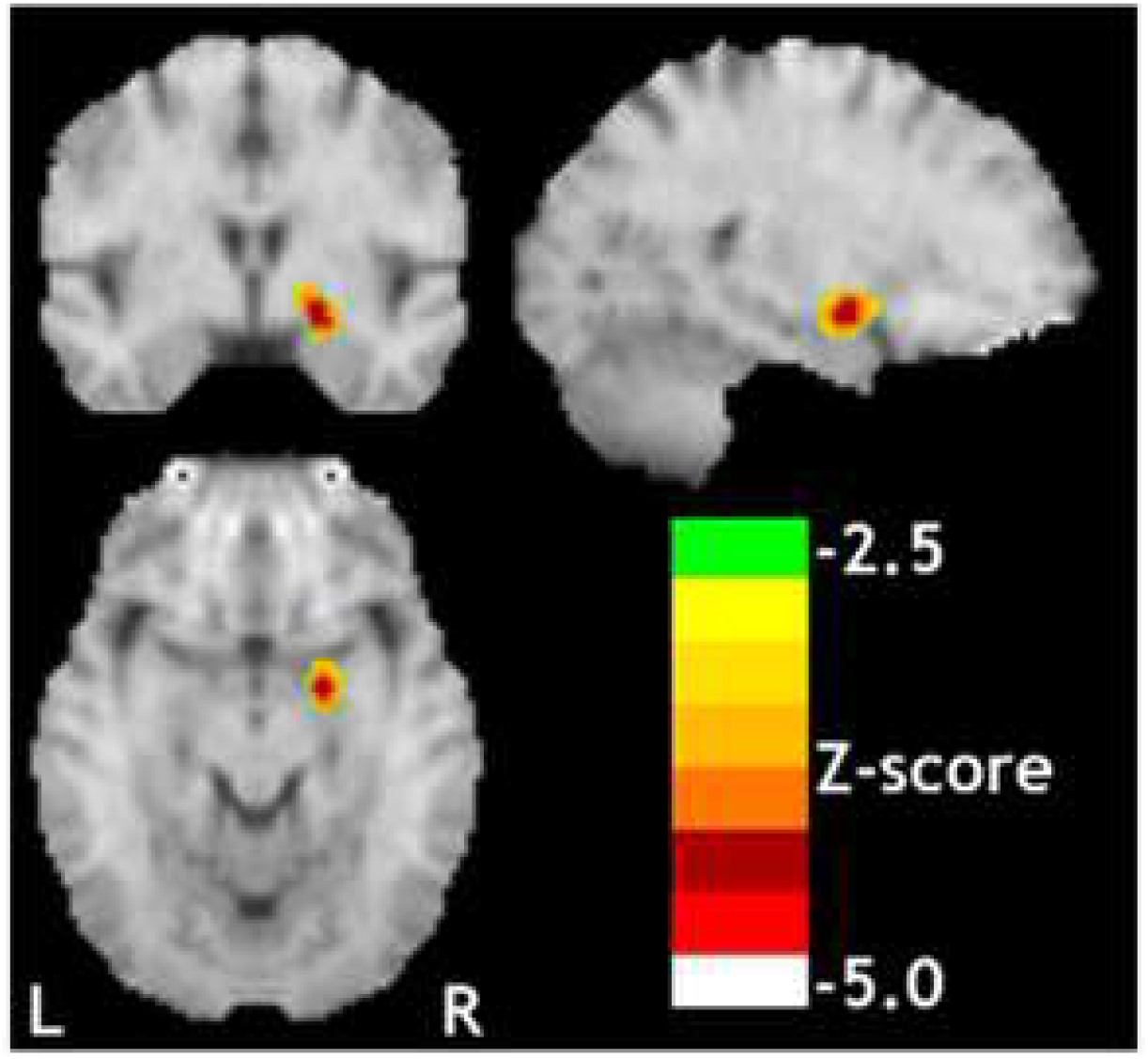
Right amygdala metabolism in responders following olanzapine/fluoxetine treatment was the only significant decrease in the whole-brain voxelwise analysis. Upper left section, coronal; lower section, transverse; upper right section, sagittal.

No studies have reported on individual metabolic patterns associated with unmedicated TRD. To explore this variability while accepting the limitation of the risk for false positive or negative responses, each TRD subject’s baseline warped FDG PET (i.e., after medication washout or baseline) was contrasted with those of a normative database with threshold set at *t* =2.0 for visualization as performed previously [42]. These individual subtractions for the nine completers are show in S5 Figure including both increases and decreases in metabolism. Examination of individual subject’s scans showed considerable heterogeneity in baseline scans with a mixture of positive, negative, or null changes in the BA25/VMPFC region. This variability suggests baseline heterogeneity in TRD or metabolic changes related to the previous history of failed treatments for TRD. If any antidepressant response occurred, a relationship to metabolic signature was not evident.

### Region of interest analyses

#### ANOVA on amygdala, hippocampus, and VMPFC

The repeated measures ANOVA indicated significant main effects of TIME (*F*(1,8) = 5.992, *p* = 0.04) and ROI (*F*(2,16) = 24.94, *p* = 0.001) without significant interaction effects (S6 Figure). This result indicates that O/F reduces activity in all three ROIs. The analysis of the correlation matrix for the ROIs reached significance only for the correlation between the left and right sgACC post-treatment (*r* = 0.82, *p* = 0.007; S2 Table). O/F tended to increase each region’s inter-hemispheric functional connectivity compared to baseline.

#### Amygdala region

Responders showed a significant reduction in mean glucose metabolism in the right amygdala ROI (*t*(3) = 3.38, *p* = 0.04), while non-responders showed no change with treatment (*t*(4) = −0.68, *p* = 0.54) (Fig 3A). There was no change in left amygdala metabolism in either group (responders: *t*(3) = 0.88, *p* = 0.45; non-responders: *t*(4) = 1.29, *p* = 0.28) (Figure 3A). Regression analyses showed no correlation between percent reduction in left amygdala metabolism and percent-reduction in MADRS scores across all subjects (Fig 3B, *r* = −0.20, *p* = 0.61). This correlation was significant in the right amygdala (Fig 3B, *r* = −0.70, *p* = 0.03). Because the hippocampus is near the amygdala and the resolution of PET in this study is low, the post hoc hippocampal region was also examined for changes in activity.

**Figure 3.**
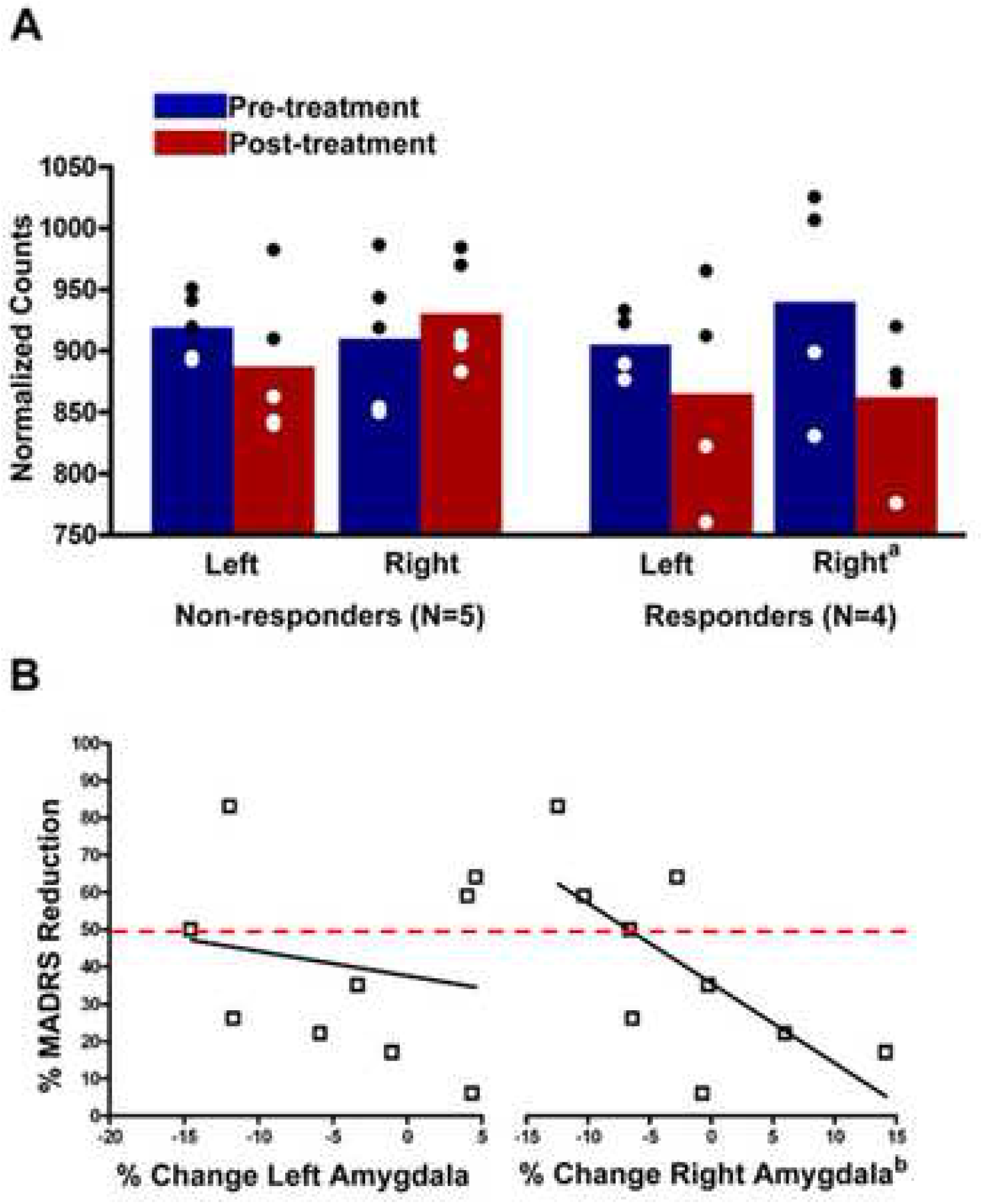
Changes in amygdala glucose uptake before and after treatment and its relationship to depression symptoms. (A) Amygdala metabolism examined separately for responders vs. non-responders in the right and left amygdala before and after treatment. ^a^ p < 0.03. (B) Change in MADRS scores and change in amygdala metabolism. Only the right amygdala showed a significant correlation between change in metabolism and change in MADRS scores. Red line identifies threshold for response.

#### Exploratory subgenual regions (BA 25 & sgACC/VMPFC)

Paired *t*-tests revealed no changes with treatment in either the left or right BA25 metabolism in either responders (left: *t*(3) = 0.05, *p* = 0.97; right: *t*(3) = 0.71, *p* = 0.53) or non-responders (left: *t*(4) = 1.13, *p* = 0.32; right: *t*(4) = 0.67, *p* = 0.54). Changes in left or right BA25 metabolism showed no correlation with changes from baseline MADRS (left: *r* = 0.29, *p* = 0.45; right: *r* = 0.15, *p* = 0.69). However, baseline right hemisphere BA25 metabolism showed a marginally significant correlation with change from baseline MADRS, whereby higher baseline metabolism predicted better response to O/F therapy (*r* = 0.68, *p* = 0.05). This correlation was not significant in the left hemisphere (*r* = 0.13, *p* = 0.74).

Paired t-tests revealed a significant decrease in right but not left VMPFC metabolism in responders (left: *t*(3) = 0.07, *p* = 0.95; right: *t*(3) = 4.18, *p* = 0.02), and no significant change in VMPFC metabolism in non-responders (left: *t*(4) = 0.81, *p* = 0.47; right: *t*(4) = 0.53, *p* = 0.62). Changes in VMPFC metabolism showed no correlation with change in MADRS (left: *r* = 0.11; *p* = 0.77; right: *r* = 0.18, *p* = 0.64). Baseline metabolism in the right, but not left, VMPFC area correlated with MADRS reduction, with higher baseline metabolism predicting better response (left: *r* = 0.37, *p* = 0.33; right: *r* = 0.84, *p* < 0.01).

## Discussion

This preliminary study found that medication-free TRD patients treated with O/F at therapeutic doses for an adequate duration showed a response-related decline in the metabolism of the right dorsal amygdala using a whole-brain, voxel-wise analysis. The dorsal amygdala in humans consists mostly of the central nucleus, the terminus of the spino-parabrachial-amygdaloid pain pathway and the principal efferent pathway for the emotional and physiological processing. Given the low resolution of the present study, caution is warranted in localization pending higher resolution techniques (e.g., higher resolution PET scanners and coregistered high resolution MRI). Several increases in metabolism surfaced also after treatment.

Two regions of interest (and one post-hoc region) based on existing literature were examined for drug-related changes in metabolism and relationship to response. Following treatment, the change in metabolism of the right amygdala correlated with the change in MADRS score; no correlation was observed for the left amygdala. Treatment did not significantly change the smaller sgACC region defined as BA25. However, a broader ROI along the VMPFC approached significance despite the small sample size.

ANOVA of ROIs in the VMPFC, amygdala, and immediately adjacent hippocampus (a post hoc exploratory ROI) indicated a main effect of treatment and ROI without significant interactions. The VMPFC showed the highest activity. O/F was associated with reduced activity in all ROIs tested. The hippocampus did not appear responsible for the deactivation seen in the dorsal amygdala. However, the hippocampus may have followed similar changes with treatment that did not reach significance; such changes have been reported previously.

Baseline metabolism in TRD may have relevance to treatment response and may, if confirmed, find utility for patient selection in treatment trials. Greater baseline metabolism in right BA25 and sgACC/VMPFC predicted better response to O/F therapy. Also, responders showed a significant decrease in metabolism following treatment in the right VMPC region which could reflect a higher initial baseline (as BA25 was included in the right VMPFC region). The higher baseline BA25 metabolism in responders to O/F may relate to 1) higher baseline resting sgACC/VMFPC blood flow seen in TRD patients when compared to controls during neuromodulation trials [10, 11]; 2) increased sgACC blood flow induced by sadness induction [59]; and 3) and increased resting blood flow in sgACC/VMFPC in healthy subjects high in negative affect [60]. Likewise, the decline in right VMPFC activity in TRD treated with O/F is analogous to the decline in resting blood flow in VMPFC in TRD responders treated with deep brain stimulation [10] as well as decreased VMPFC metabolism with chronic vagus nerve stimulation [18, 56].

Strengths of the study include the recruitment of severely ill TRD patients, medication washout before treatment, exploratory examination of individual TRD patients, and full course of treatment with O/F to maximal tolerated doses. These preliminary data suggest heterogeneity in metabolic signatures in TRD. If replicated, this heterogeneity requires consideration in the design of trials for TRD using medications or devices. However, the changes observed with treatment appear broadly like those reported in treatment-responsive major depression with other antidepressant treatments: decreased amygdala and VMPFC metabolism.

Limitations of this study include comorbidities, prior heterogeneous treatments during past failed trials, significant participant dropout, small final sample size, lack of a placebo, low PET resolution, and confounding of depression with anxiety measures. Patient dropouts and comorbidities could limit generalizability. However, most TRD patients do have comorbidity [5-8]. A larger replication sample could address whether major comorbidities represent a covariate of interest in the response as suggested by this study—likewise for dropouts. Future studies will need to account for the large dropout during washout. The limited resolution of PET places some ambiguity in the precise determination of amygdala activity. However, the latest scanners with resolution of 3 mm full width at half maximum will improve localization in future studies. Of note, although the ROI analysis of the right amygdala confirmed significant deactivation after treatment that correlated with clinical response, the focus was only partially resolved from other nearby regions which showed a similar response pattern (e.g., hippocampus). Depression and anxiety are both prominent symptoms in TRD, and antidepressants/atypical neuroleptics often decrease both depression and anxiety scores. In this regard, HAMD and HAMA scores are inter-correlated [61]. Anxiety scores measured in this study in responders before and after treatments did not differ. Therefore, the decrease in amygdala activity does not likely reflect changes in anxiety, but rather depression. The severity of illness and risk of suicide precluded a placebo control. However, other studies using FDG PET and placebos in test-retest designs suggest high consistency in normalized regional activity and only small changes [62-64].

## Conclusions

TRD patients show considerable baseline metabolic heterogeneity following medication washout. Whether this heterogeneity arises from differing disease pathologies or from effects of past treatments remains unclear. As reported for other antidepressant therapies in treatment-responsive depression, decline in amygdala and VMPFC activity surfaced here with O/F treatment. Furthermore, decreased metabolism in the right amygdala with treatment correlated with improvement in depression following O/F treatment.

## Acknowledgments

This work was supported through an Investigator-Initiated Grant from Eli Lilly & Company and the Department of Veterans Affairs (I01CX000501). A fixed dose of olanzapine and fluoxetine combination with trade name Symbyax has been approved by the USA Food and Drug Administration with indications for TRD and bipolar I depression. The funders had no role in study design, data collection and analysis, decision to publish, or preparation of the manuscript. We thank Hemant Shah for assistance in data collection and thank the volunteers for their patience and perseverance.

## Authors’ Contributions

JVP designed the study, secured funding, participated in data collection and analyses, and wrote/edited the final manuscript. SAS designed the study, assisted in securing funding, participated in data collection and reviewed and edited the final manuscript. GS analyzed data, provided figures, wrote the initial manuscript, and reviewed and edited the final manuscript. JTL analyzed data, contributed software, curated data, provided figures, and edited the final manuscript. DA, BR, FSA provided clinical care and recruitment of the patients, gathered clinical data, and edited the final version of the manuscript.

## Supporting Information

**S1 Figure. Weights before and after O/F combination**.

**S2 Figure. Brain glucose uptake in non-responders: Post- minus Pre- O/F treatment**.

Stereotactically normalized. Image left is right side of brain. AC-PC plane 0 mm. Color scale shows Z-scores with threshold Z = ±3.3.

**S3 Figure. Brain glucose uptake in responders: Post- minus Pre- O/F treatment**

Stereotactically normalized. Image left is right side of brain. AC-PC plane 0 mm. Color scale shows Z-scores with threshold at Z = ±3.3.

**S4 Figure. Brain glucose uptake for all subjects: Post- minus Pre- O/F treatment**.

Stereotactically normalized. Image left is right side of brain. AC-PC plane 0 mm. Color scale shows Z-scores with threshold at Z = ±3.3.

**S5 Figure. Differences in resting brain glucose uptake between individual subjects (N = 9) at baseline (after washout) and a normative data set (N = 30)**.

For visualizing individual metabolic fingerprints of all nine subjects, the threshold was set at t = 2.0 that is the usual threshold used for studying change in individuals [42]. Each subject is represented by a study number (e.g., pL0009). Age regression was used to match individual subject’s age to that of the normative group. R, right; L, left, A, anterior; P, posterior. The patterns are heterogenous. For example, some individuals have sgACC/VMPFC hypoactive, hyperactivity, or no change.

**S6 Figure. Main effects of TIME and ROI on glucose uptake**.

**S1 Table. Individual subject’s weight, depression scores, and anxiety ratings**.

**S2 Table. Correlation matrix for metabolism in bilateral ROIs**.

Green cells below diagonal are for Pre-treatment; blue cells above diagonal are for Post-treatment. R, right; L, left; Hippo, hippocampus; sgACC, subgenual anterior cingulate/VMPFC. ^†^ p < 0.007

