## Supplementary info file for "A preliminary study of resting brain metabolism in treatment-resistant depression before and after treatment with olanzapine-fluoxetine combination"

#### Supporting Information

**S1 Figure. Weights before and after O/F combination.**

**S2 Figure. Brain glucose uptake in non-responders: Post- minus Pre- O/F treatment.**

Stereotactically normalized. Image left is right side of brain. AC-PC plane 0 mm. Color scale shows Z-scores with threshold Z = ±3.3.

**S3 Figure. Brain glucose uptake in responders: Post- minus Pre- O/F treatment**

Stereotactically normalized. Image left is right side of brain. AC-PC plane 0 mm. Color scale shows Z-scores with threshold at Z = ±3.3.

**S4 Figure. Brain glucose uptake for all subjects: Post- minus Pre- O/F treatment.**

Stereotactically normalized. Image left is right side of brain. AC-PC plane 0 mm. Color scale shows Z-scores with threshold at Z = ±3.3.

**S5 Figure. Differences in resting brain glucose uptake between individual subjects (N = 9) at baseline (after washout) and a normative data set (N = 30).**

For visualizing individual metabolic fingerprints of all nine subjects, the threshold was set at t = 2.0 that is the usual threshold used for studying change in individuals [42]. Each subject is represented by a study number (e.g., pL0009). Age regression was used to match individual subject's age to that of the normative group. R, right; L, left, A, anterior; P, posterior. The patterns are heterogenous. For example, some individuals have sgACC/VMPFC hypoactive, hyperactivity, or no change.

**S6 Figure. Main effects of TIME and ROI on glucose uptake.**

**S1 Table. Individual subject's weight, depression scores, and anxiety ratings.**

**S2 Table. Correlation matrix for metabolism in bilateral ROIs.**

Green cells below diagonal are for Pre-treatment; blue cells above diagonal are for Post-treatment. R, right; L, left; Hippo, hippocampus; sgACC, subgenual anterior cingulate/VMPFC. [†] $p < 0.007$

S1 Figure.

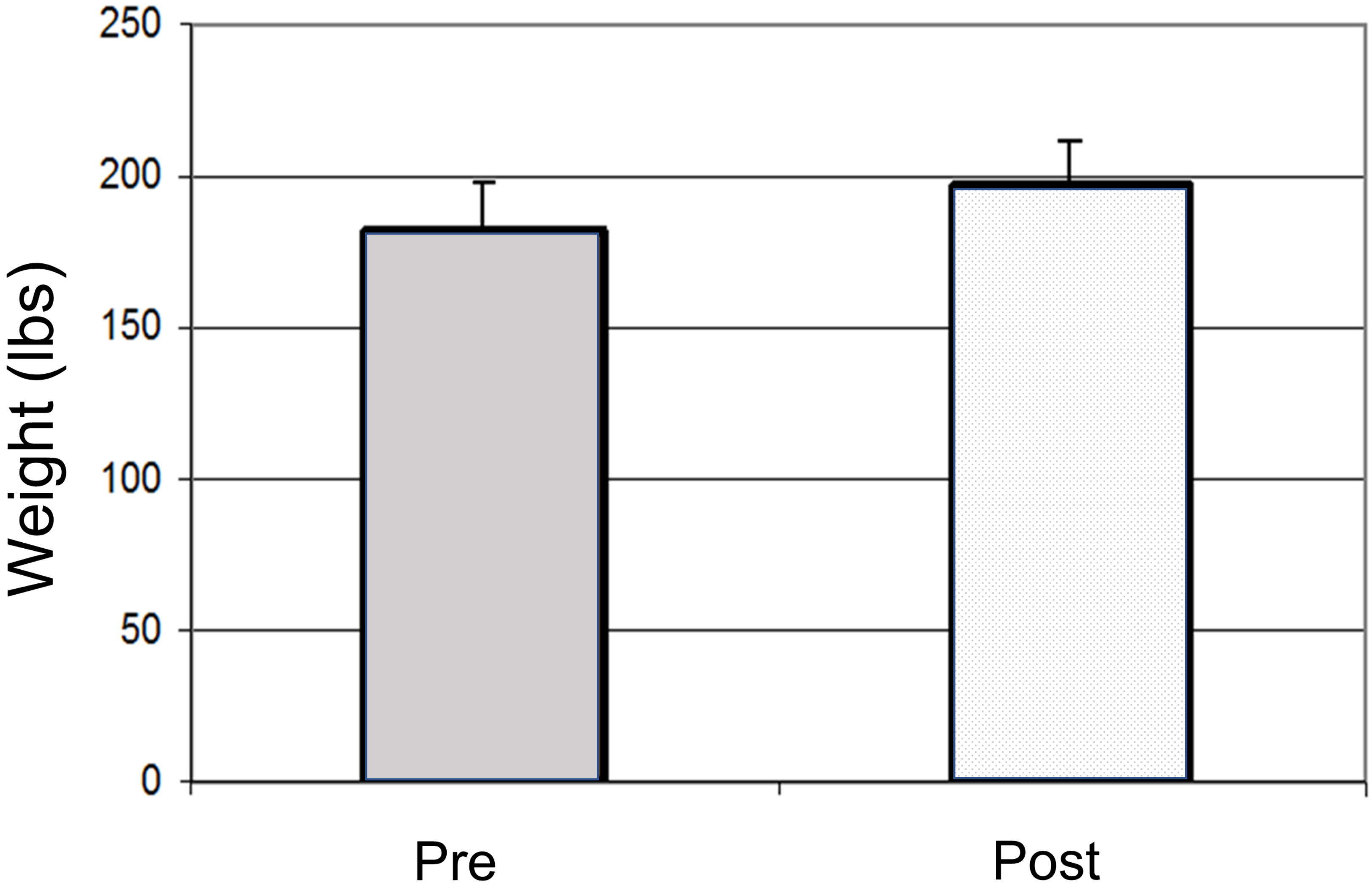

### S2 Figure.

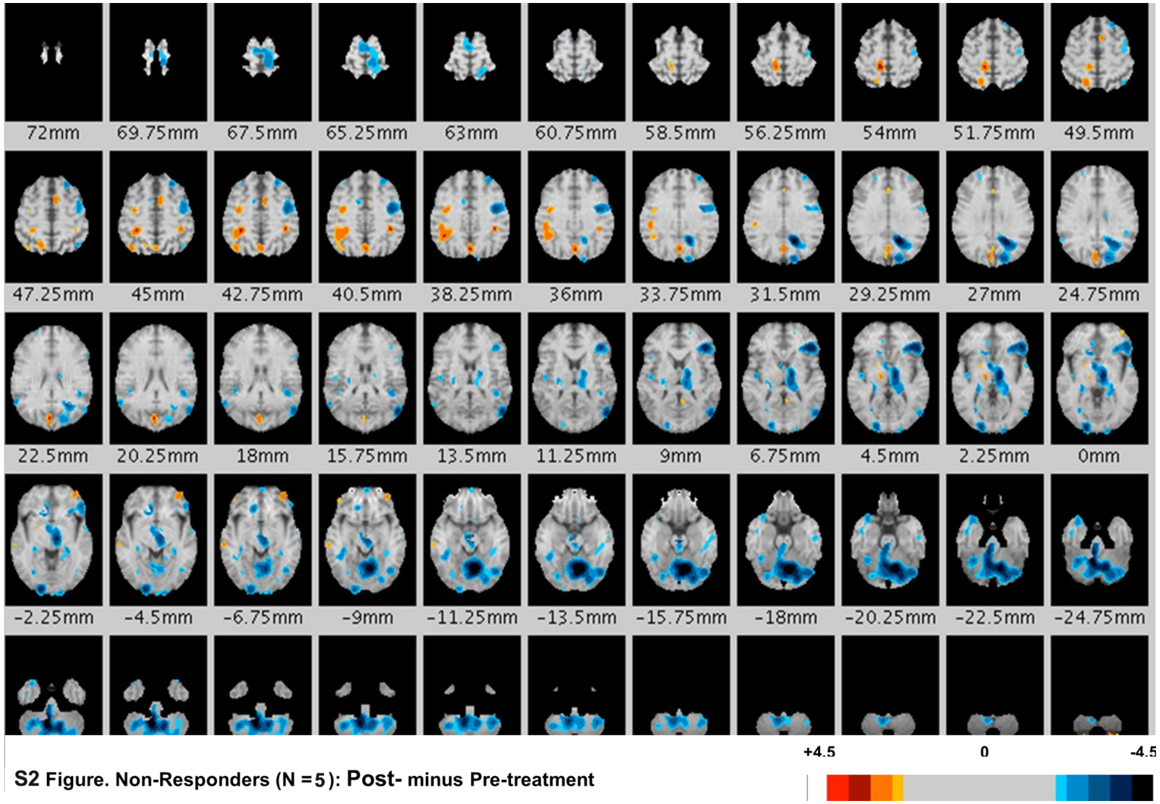


**S2 Figure. Non-Responders (N = 5): Post- minus Pre-treatment**

### S3 Figure.

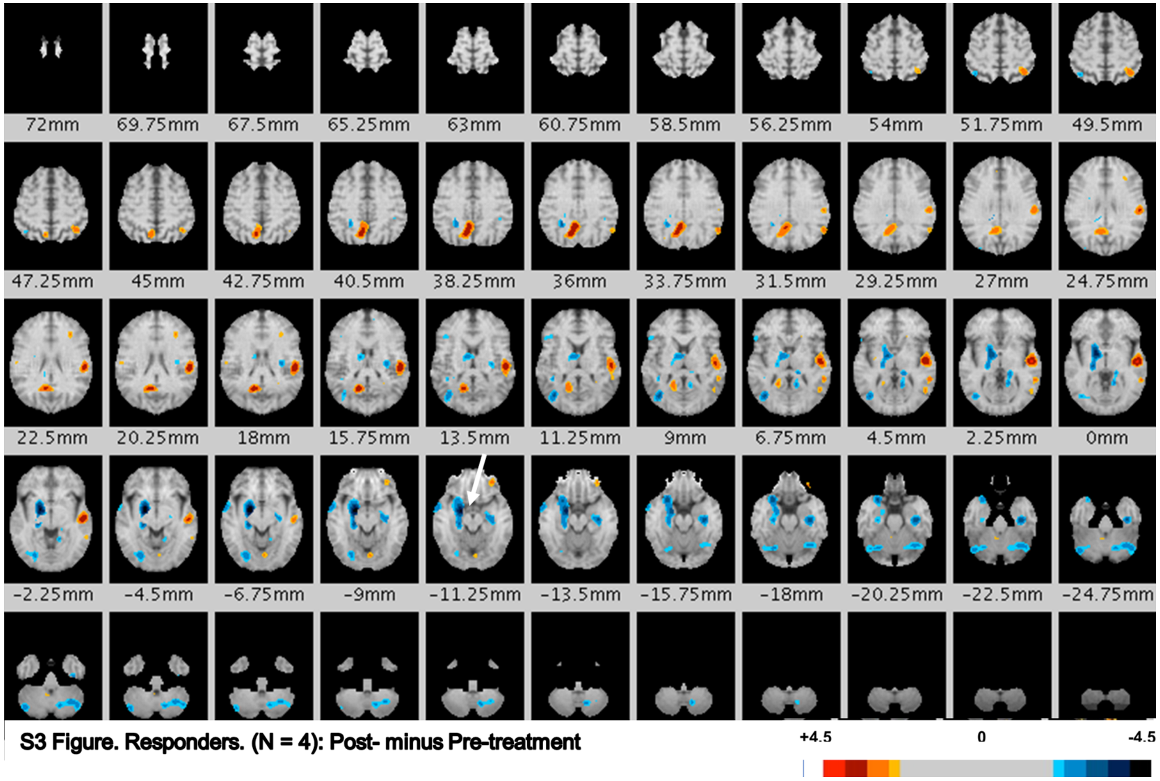


**S3 Figure. Responders. (N = 4): Post- minus Pre-treatment**

### S4 Figure.

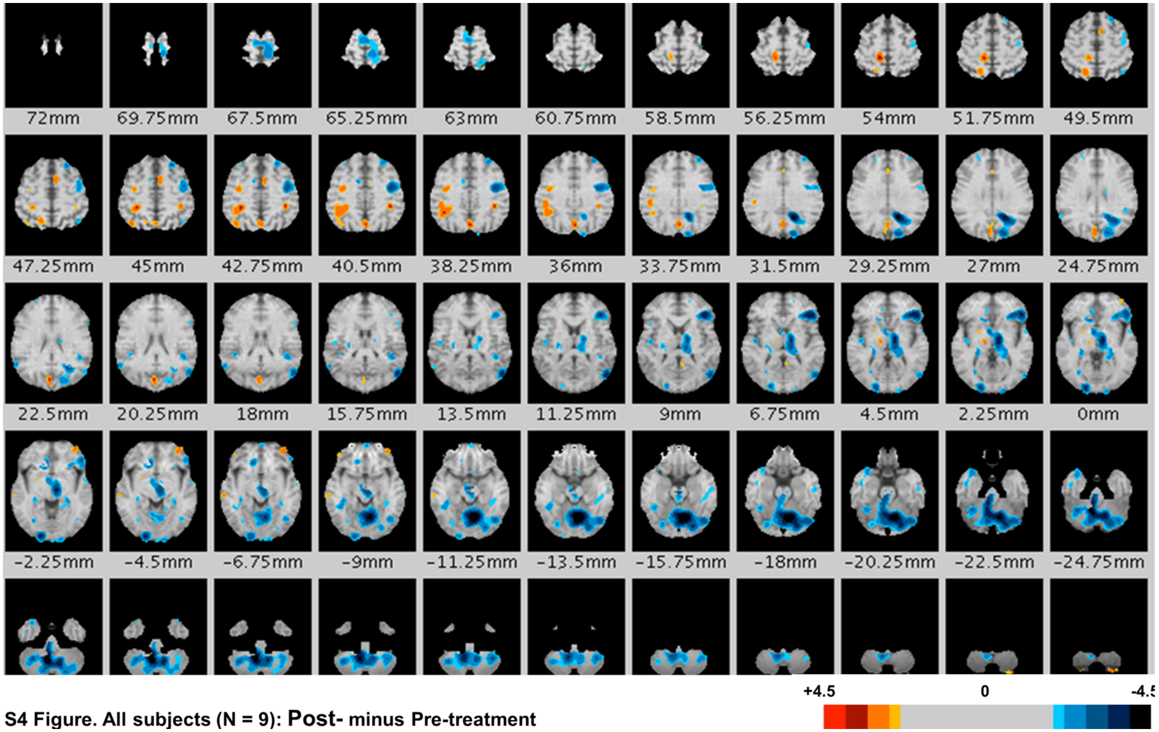


**S4 Figure. All subjects (N = 9): Post- minus Pre-treatment**

S5 Figure. Please see S5 Figure following S6 Figure (below) that contains each individual's change in resting glucose uptake contrasted with that of normal subjects all stereotactically normalized and regressed for age.

S6 Figure.

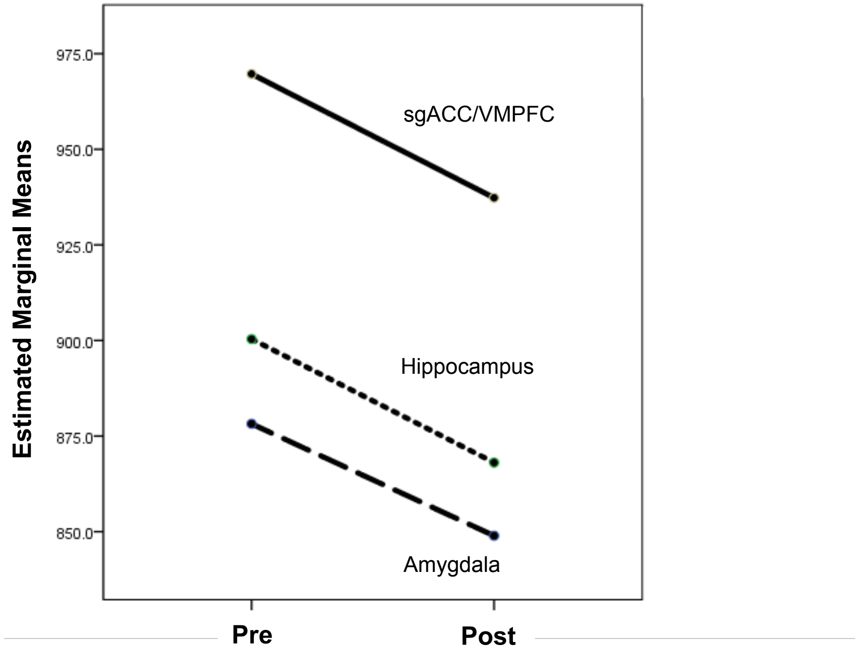
Estimated Marginal Means
975.0
950.0
925.0
900.0
875.0
850.0
sgACC/VMPFC
Hippocampus
Amygdala
Pre
Post

### S1 Table.

**S1 Table. Pre- vs. Post- Weights, depression scores, and anxiety scores.**

| Scan # pL | Age (years) | Weight pre lb | Weight post lb | MADRS pre | MADRS post | HAMA pre | HAMA post |
|---|---|---|---|---|---|---|---|
| 0020/0021 | 47 | 100 | 132 | 33 | 31 | 25 | 16 |
| 0009/0011 | 44 | 279 | 288 | 29 | 24 | 25 | 19 |
| 0026/0027 | 52 | 163 | 176 | 37 | 29 | 15 | 8 |
| 0028/0029 | 36 | 217 | 235 | 27 | 20 | 9 | 7 |
| 0030/0031 | 62 | 185 | 188 | 29 | 12 | 16 | 16 |
| 0059/0069 | 54 | 198 | 205 | 24 | 4 | 17 | 11 |
| 0071/0072 | 61 | 187 | 210 | 31 | 20 | 12 | 6 |
| 0079/0087 | 27 | 162 | 173 | 38 | 19 | 29 | 15 |
| 0089/0095 | 41 | 142 | 164 | 28 | 10 | 14 | 10 |

lb, pounds; MADRS, Madras Asberg Depression Rating Scale; HAMA, Hamilton Anxiety Scale.

### S2 Table.

**S2 Table. Correlation matrix for metabolism in ROIs.**

| | R amygdala | L amygdala | R Hippo | L Hippo | R sgACC | L sgACC |
|---|---|---|---|---|---|---|
| R amygdala | 1 | 0.611 | 0.604 | 0.583 | 0.074 | -0.008 |
| L amygdala | 0.011 | 1 | 0.241 | 0.232 | 0.052 | -0.040 |
| R Hippo | 0.595 | 0.173 | 1 | 0.449 | 0.244 | -0.031 |
| L Hippo | 0.210 | -0.033 | -0.115 | 1 | 0.125 | -0.296 |
| R sgACC | 0.360 | -0.341 | 0.425 | 0.004 | 1 | 0.821† |
| L sgACC | 0.527 | 0.502 | 0.447 | 0.069 | 0.283 | 1 |

Green cells below diagonal are for Pre-treatment; blue cells above diagonal are for Post-treatment. R, right; L, left; Hippo, hippocampus; sgACC, subgenual anterior cingulate/VMPFC.

† $p < 0.007$

S 5 Figures all subjects

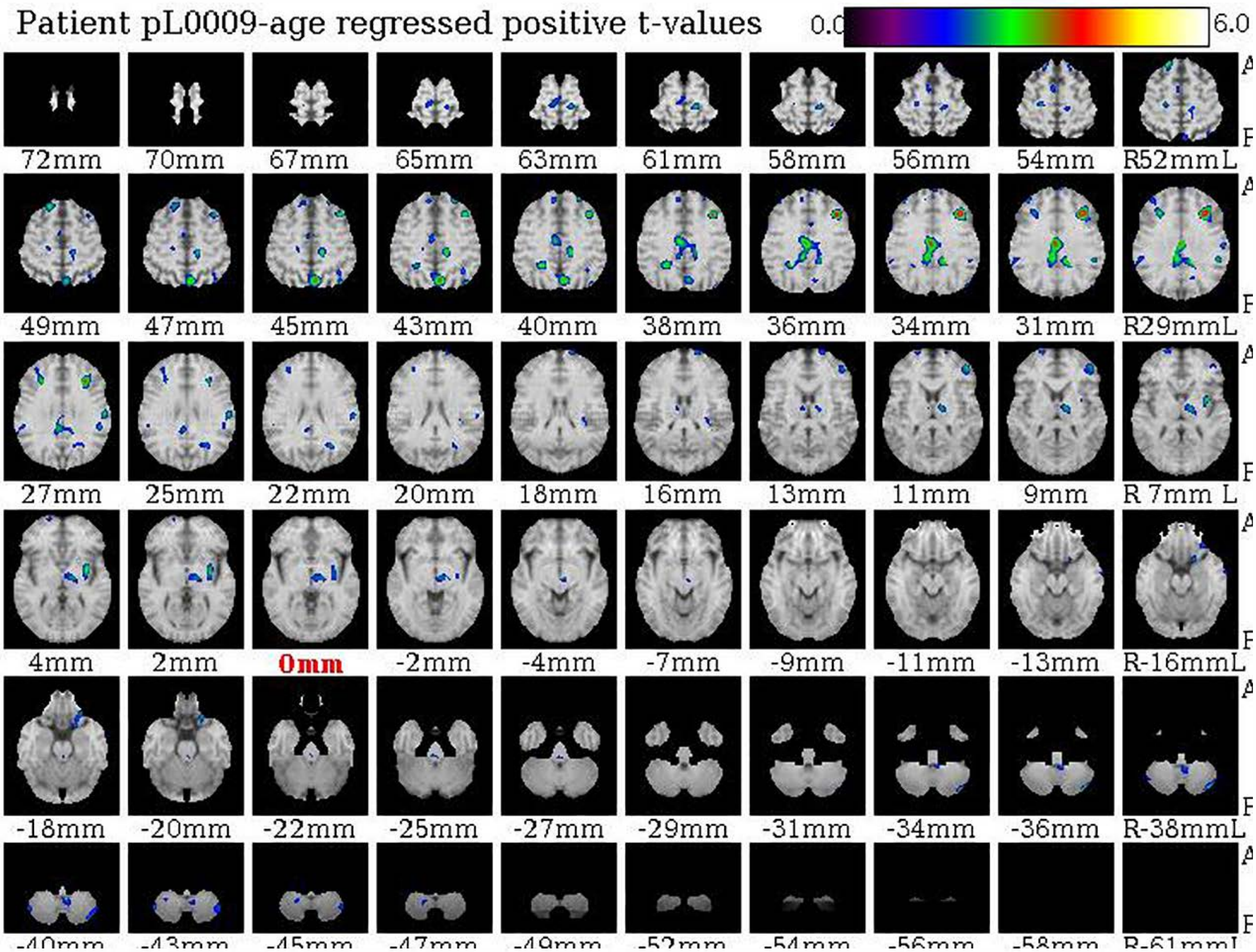

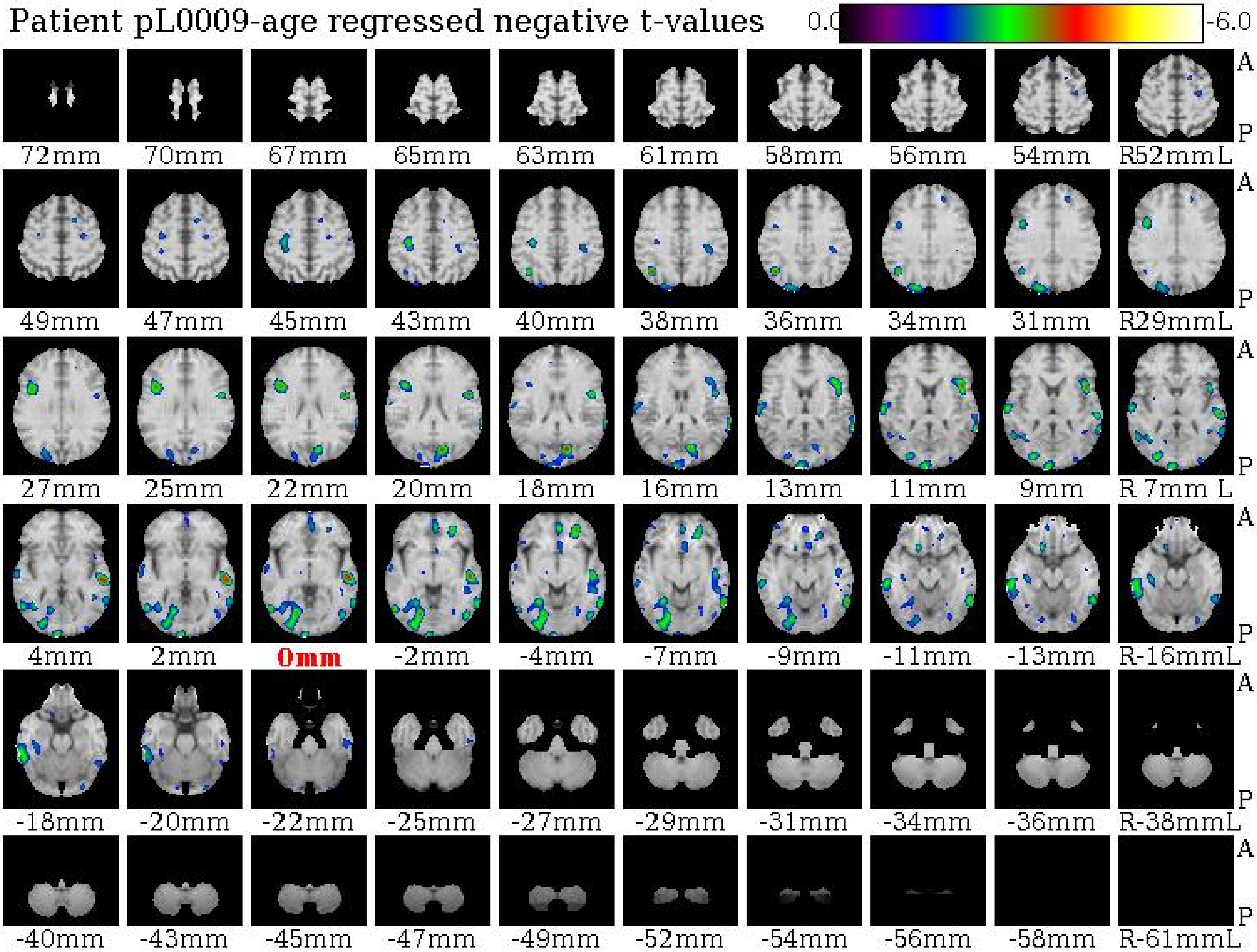

Patient pL0009-age regressed negative t-values
0.0
-6.0
A
P
72mm
70mm
67mm
65mm
63mm
61mm
58mm
56mm
54mm
R52mmL
49mm
47mm
45mm
43mm
40mm
38mm
36mm
34mm
31mm
R29mmL
27mm
25mm
22mm
20mm
18mm
16mm
13mm
11mm
9mm
R 7mm L
4mm
2mm
0mm
-2mm
-4mm
-7mm
-9mm
-11mm
-13mm
R-16mmL
-18mm
-20mm
-22mm
-25mm
-27mm
-29mm
-31mm
-34mm
-36mm
R-38mmL
-40mm
-43mm
-45mm
-47mm
-49mm
-52mm
-54mm
-56mm
-58mm
R-61mmL

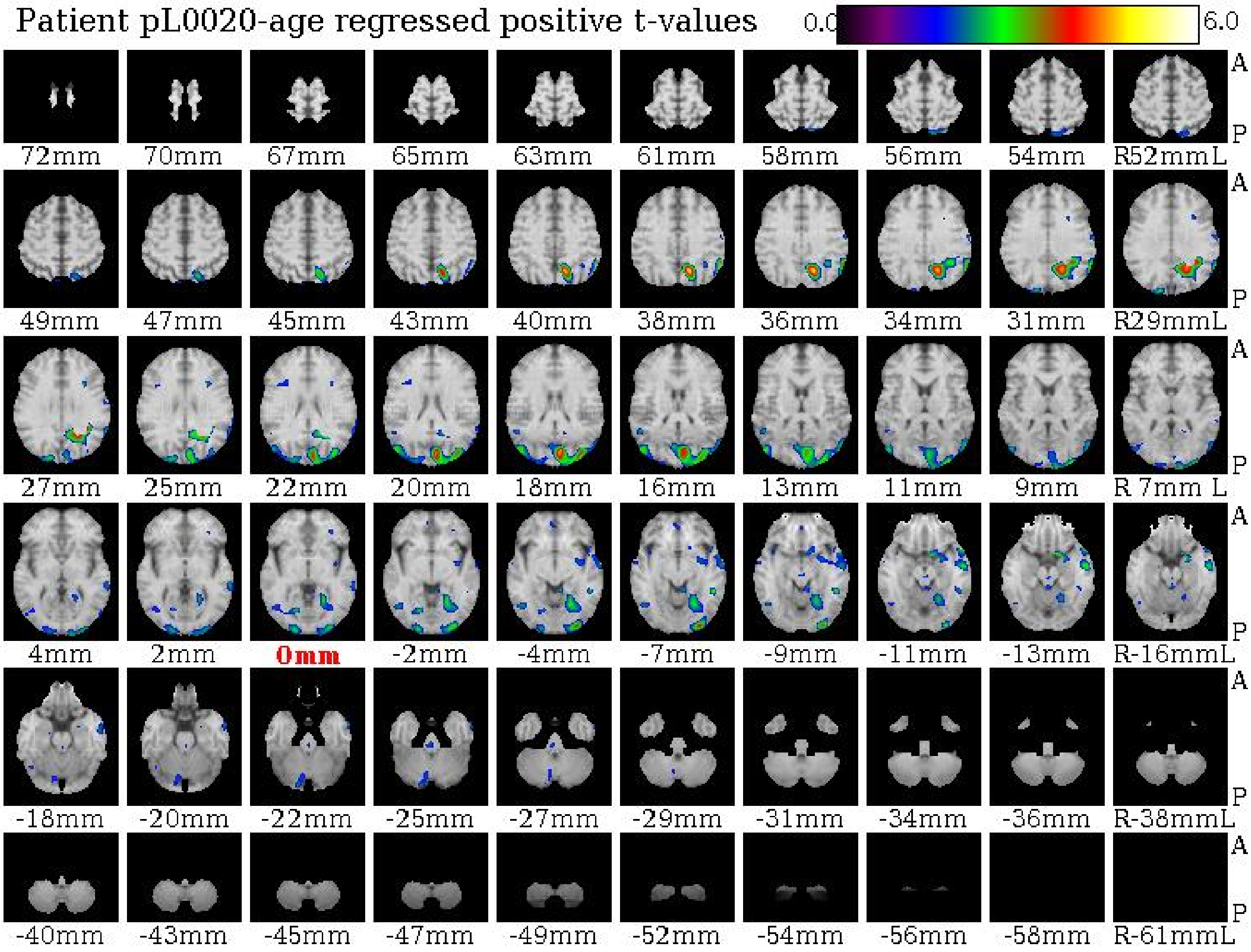

Patient pL0020-age regressed positive t-values
0.0
6.0
A
P
72mm
70mm
67mm
65mm
63mm
61mm
58mm
56mm
54mm
R52mmL
49mm
47mm
45mm
43mm
40mm
38mm
36mm
34mm
31mm
R29mmL
27mm
25mm
22mm
20mm
18mm
16mm
13mm
11mm
9mm
R 7mm L
4mm
2mm
0mm
-2mm
-4mm
-7mm
-9mm
-11mm
-13mm
R-16mmL
-18mm
-20mm
-22mm
-25mm
-27mm
-29mm
-31mm
-34mm
-36mm
R-38mmL
-40mm
-43mm
-45mm
-47mm
-49mm
-52mm
-54mm
-56mm
-58mm
R-61mmL

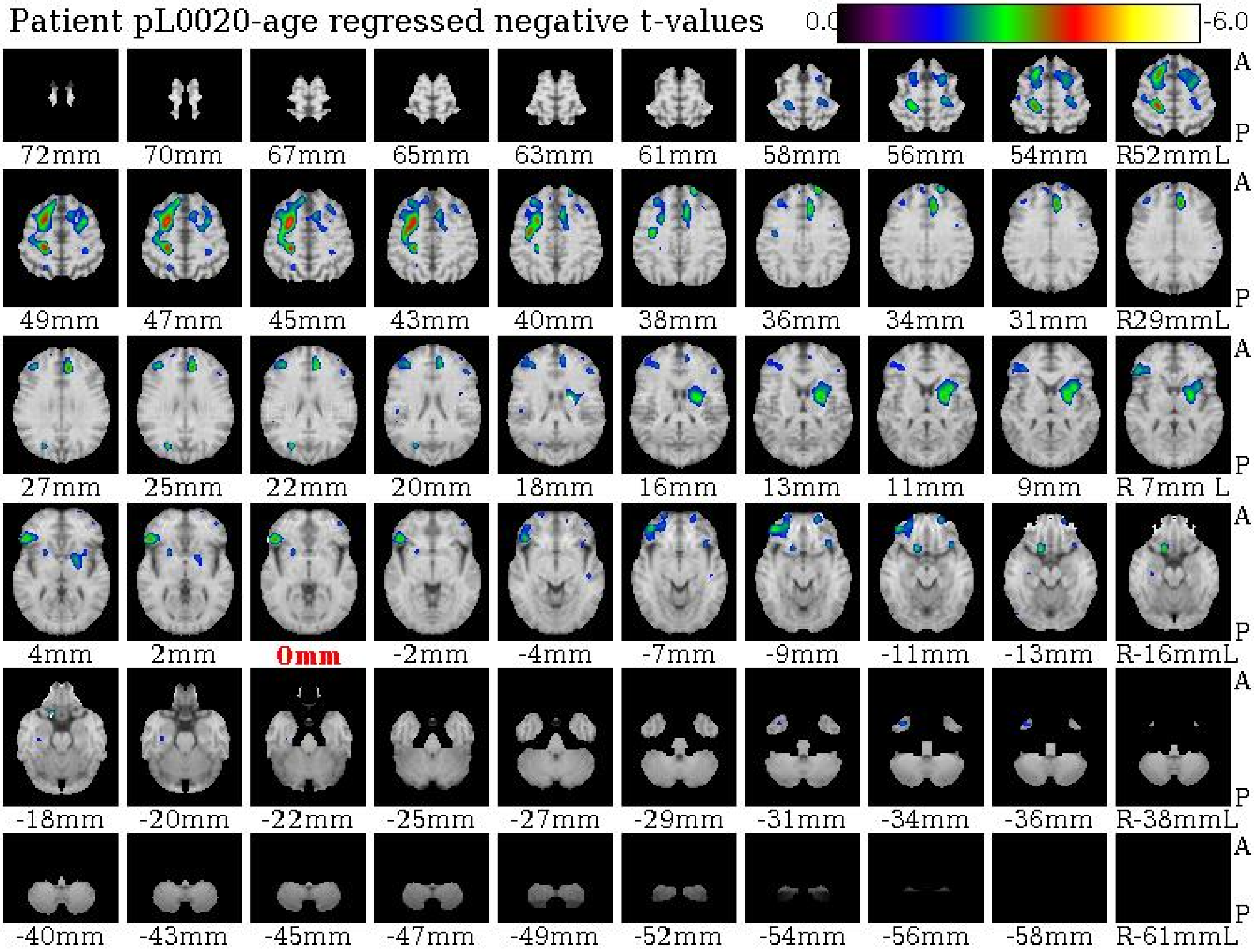

Patient pL0020-age regressed negative t-values
0.0
-6.0
72mm
70mm
67mm
65mm
63mm
61mm
58mm
56mm
54mm
R52mmL
49mm
47mm
45mm
43mm
40mm
38mm
36mm
34mm
31mm
R29mmL
27mm
25mm
22mm
20mm
18mm
16mm
13mm
11mm
9mm
R 7mm L
4mm
2mm
0mm
-2mm
-4mm
-7mm
-9mm
-11mm
-13mm
R-16mmL
-18mm
-20mm
-22mm
-25mm
-27mm
-29mm
-31mm
-34mm
-36mm
R-38mmL
-40mm
-43mm
-45mm
-47mm
-49mm
-52mm
-54mm
-56mm
-58mm
R-61mmL
A
P

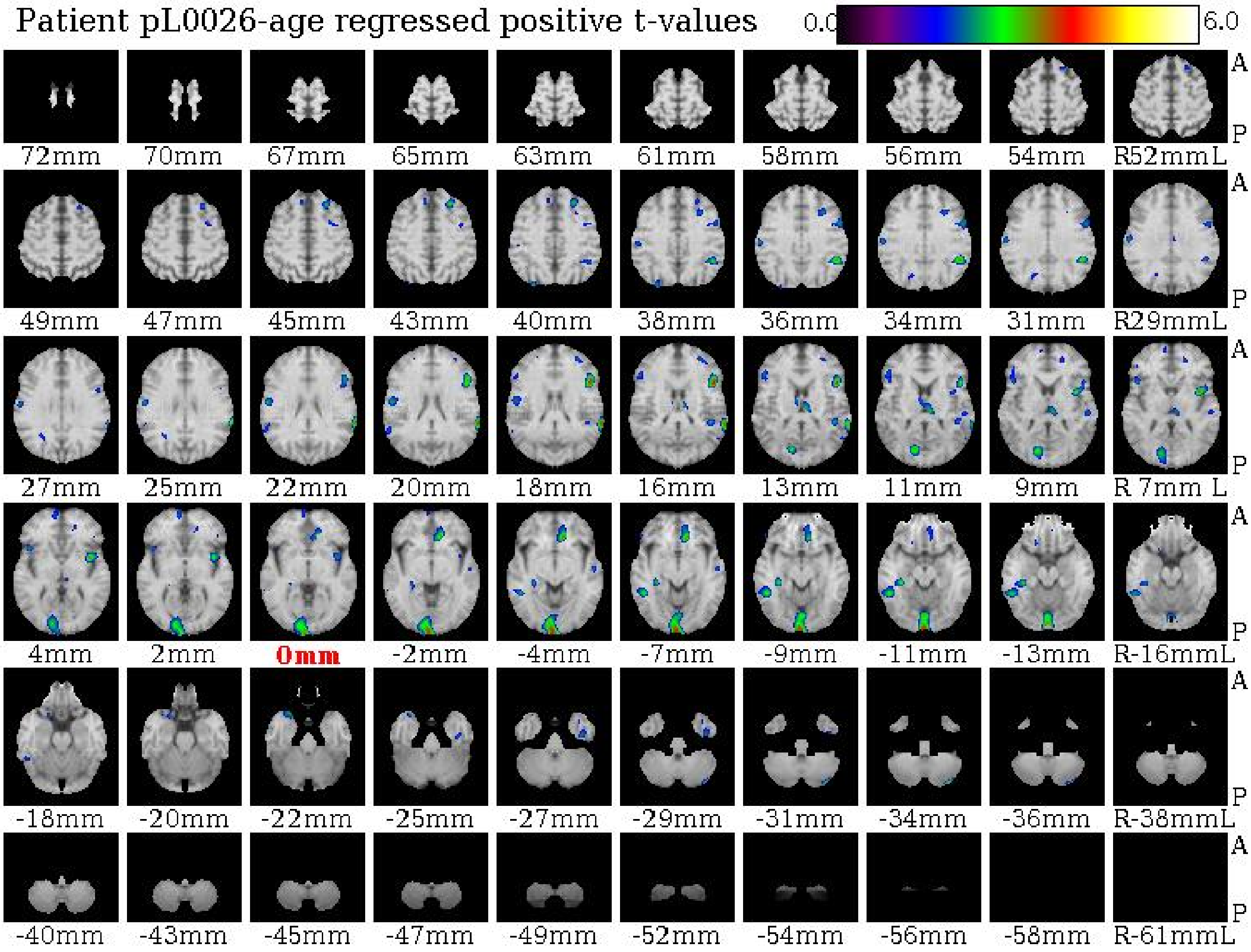
Patient pL0026-age regressed positive t-values
0.0
6.0
A
P
72mm
70mm
67mm
65mm
63mm
61mm
58mm
56mm
54mm
R52mmL
49mm
47mm
45mm
43mm
40mm
38mm
36mm
34mm
31mm
R29mmL
27mm
25mm
22mm
20mm
18mm
16mm
13mm
11mm
9mm
R 7mm L
4mm
2mm
0mm
-2mm
-4mm
-7mm
-9mm
-11mm
-13mm
R-16mmL
-18mm
-20mm
-22mm
-25mm
-27mm
-29mm
-31mm
-34mm
-36mm
R-38mmL
-40mm
-43mm
-45mm
-47mm
-49mm
-52mm
-54mm
-56mm
-58mm
R-61mmL

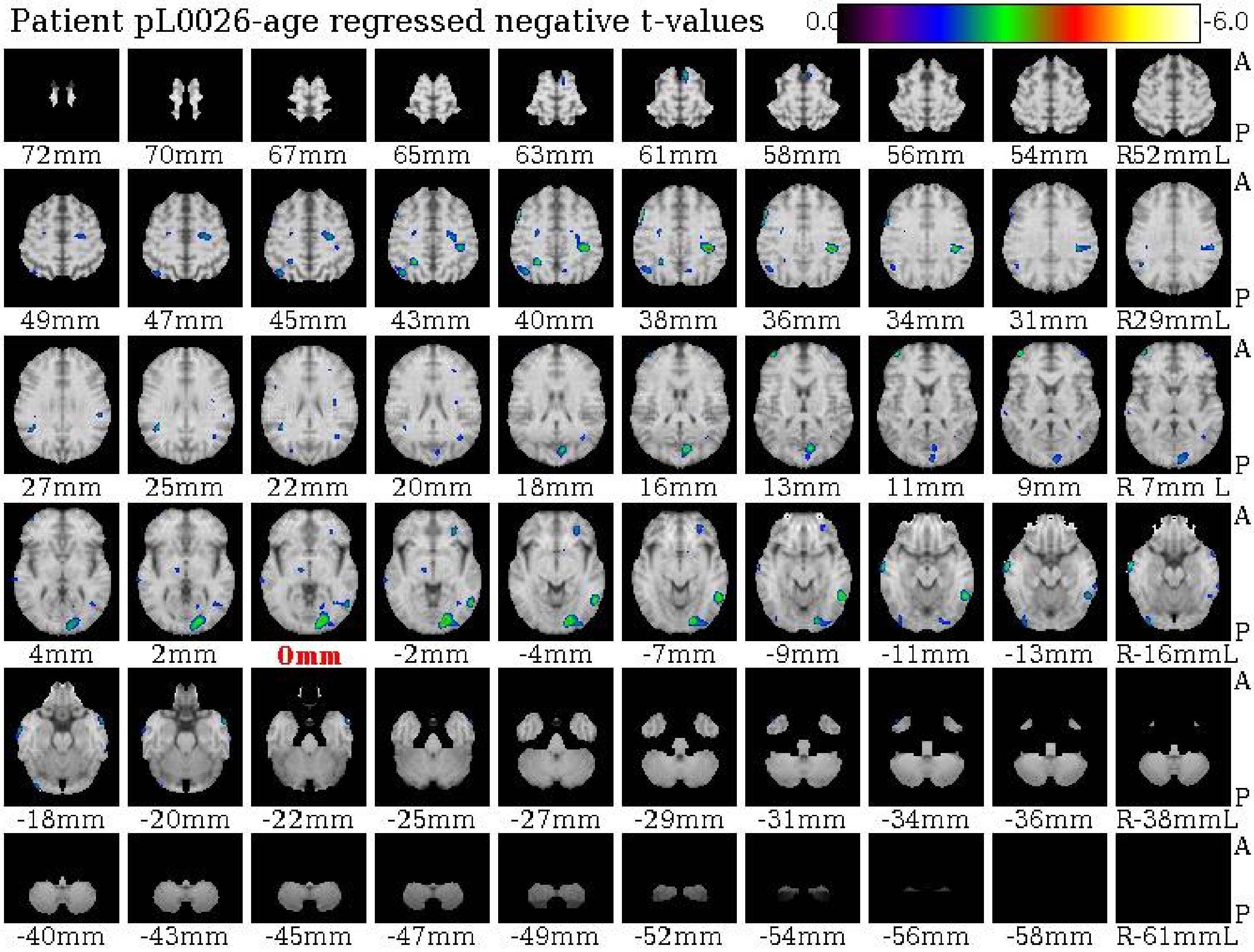

Patient pL0026-age regressed negative t-values
0.0
-6.0
A
P
72mm 70mm 67mm 65mm 63mm 61mm 58mm 56mm 54mm R52mmL
49mm 47mm 45mm 43mm 40mm 38mm 36mm 34mm 31mm R29mmL
27mm 25mm 22mm 20mm 18mm 16mm 13mm 11mm 9mm R 7mm L
4mm 2mm 0mm -2mm -4mm -7mm -9mm -11mm -13mm R-16mmL
-18mm -20mm -22mm -25mm -27mm -29mm -31mm -34mm -36mm R-38mmL
-40mm -43mm -45mm -47mm -49mm -52mm -54mm -56mm -58mm R-61mmL

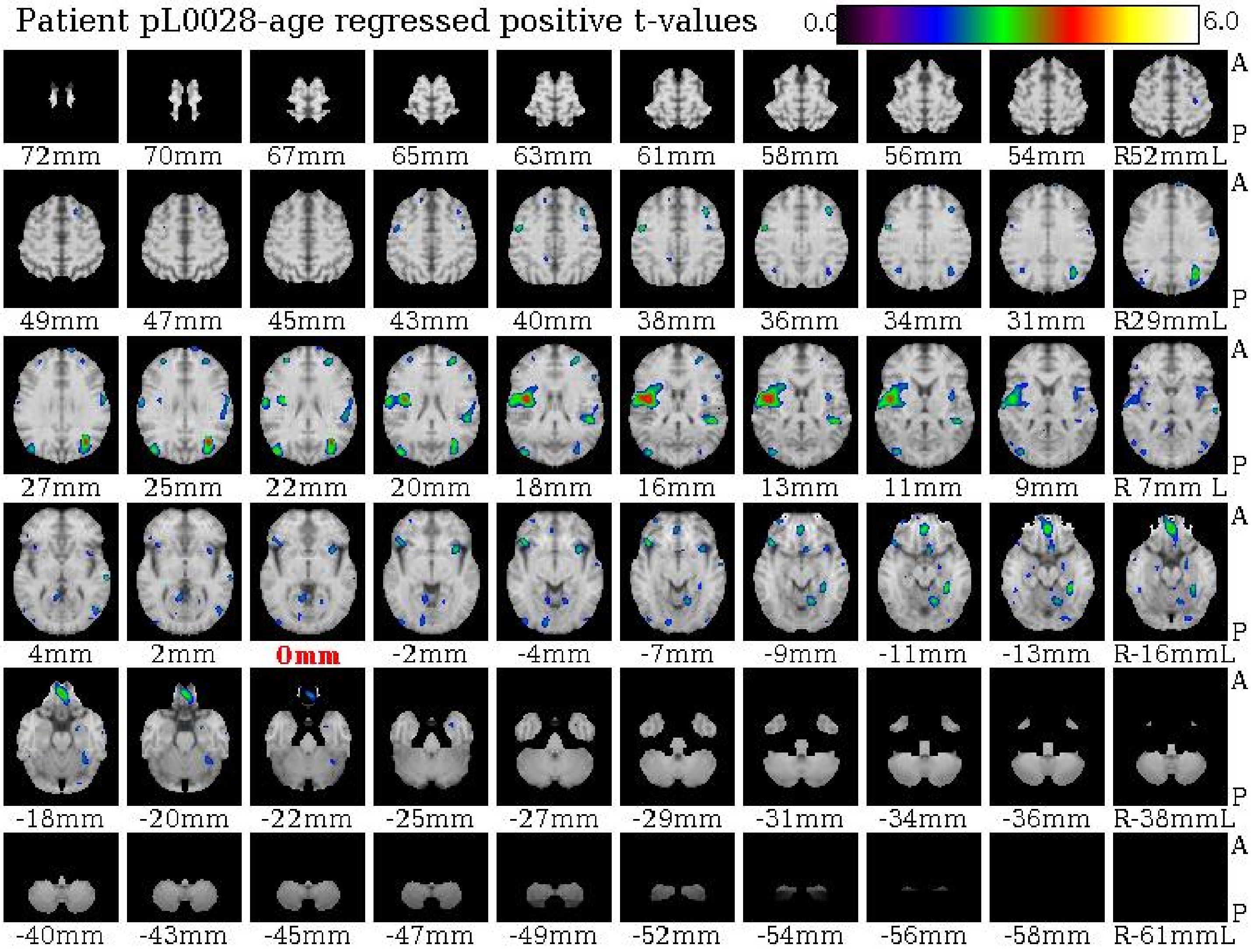
Patient pL0028-age regressed positive t-values
0.0
6.0
A
P
72mm
70mm
67mm
65mm
63mm
61mm
58mm
56mm
54mm
R52mmL
49mm
47mm
45mm
43mm
40mm
38mm
36mm
34mm
31mm
R29mmL
27mm
25mm
22mm
20mm
18mm
16mm
13mm
11mm
9mm
R 7mm L
4mm
2mm
0mm
-2mm
-4mm
-7mm
-9mm
-11mm
-13mm
R-16mmL
-18mm
-20mm
-22mm
-25mm
-27mm
-29mm
-31mm
-34mm
-36mm
R-38mmL
-40mm
-43mm
-45mm
-47mm
-49mm
-52mm
-54mm
-56mm
-58mm
R-61mmL

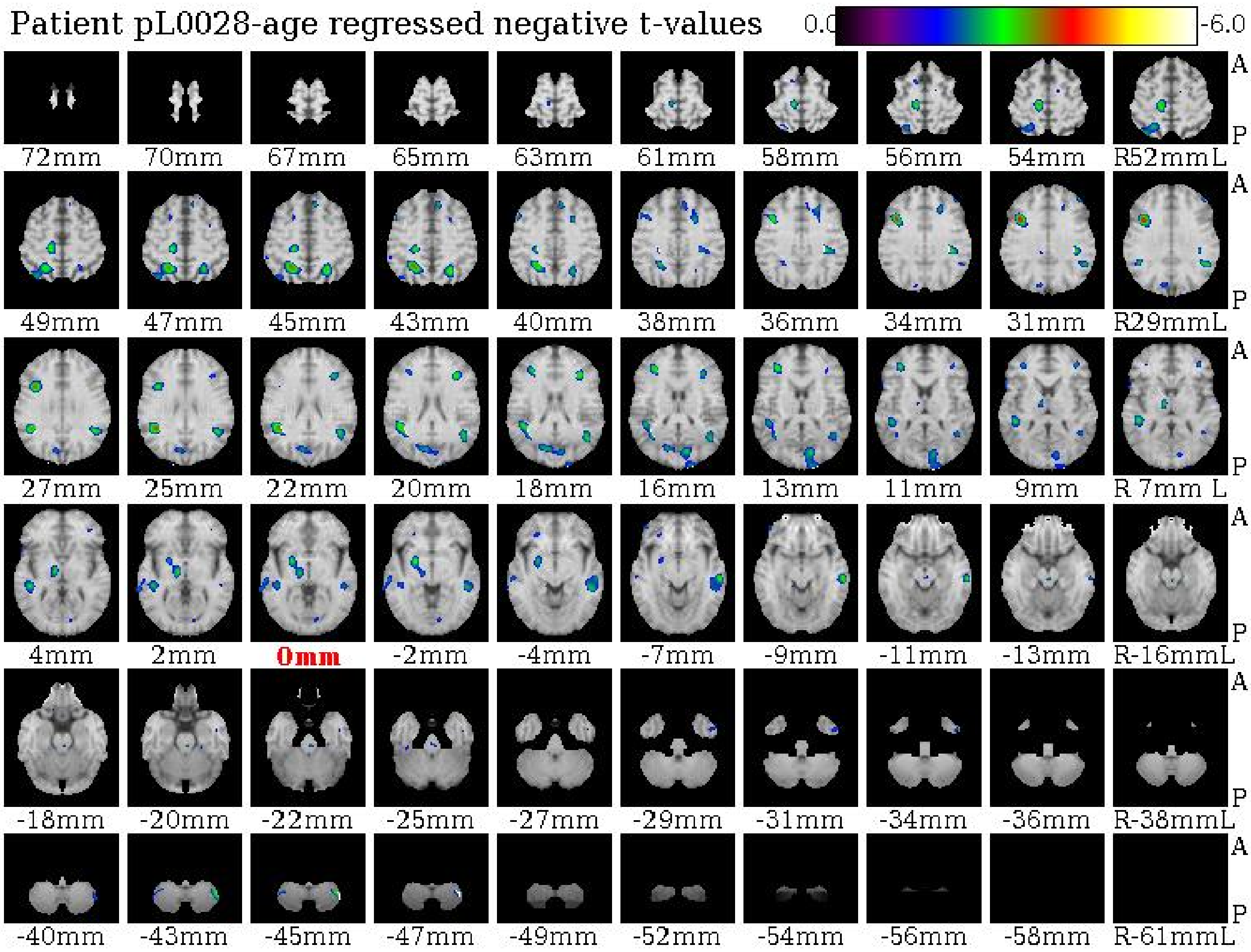
Patient pL0028-age regressed negative t-values
0.0
-6.0
72mm
70mm
67mm
65mm
63mm
61mm
58mm
56mm
54mm
R52mmL
49mm
47mm
45mm
43mm
40mm
38mm
36mm
34mm
31mm
R29mmL
27mm
25mm
22mm
20mm
18mm
16mm
13mm
11mm
9mm
R 7mm L
4mm
2mm
0mm
-2mm
-4mm
-7mm
-9mm
-11mm
-13mm
R-16mmL
-18mm
-20mm
-22mm
-25mm
-27mm
-29mm
-31mm
-34mm
-36mm
R-38mmL
-40mm
-43mm
-45mm
-47mm
-49mm
-52mm
-54mm
-56mm
-58mm
R-61mmL
A
P

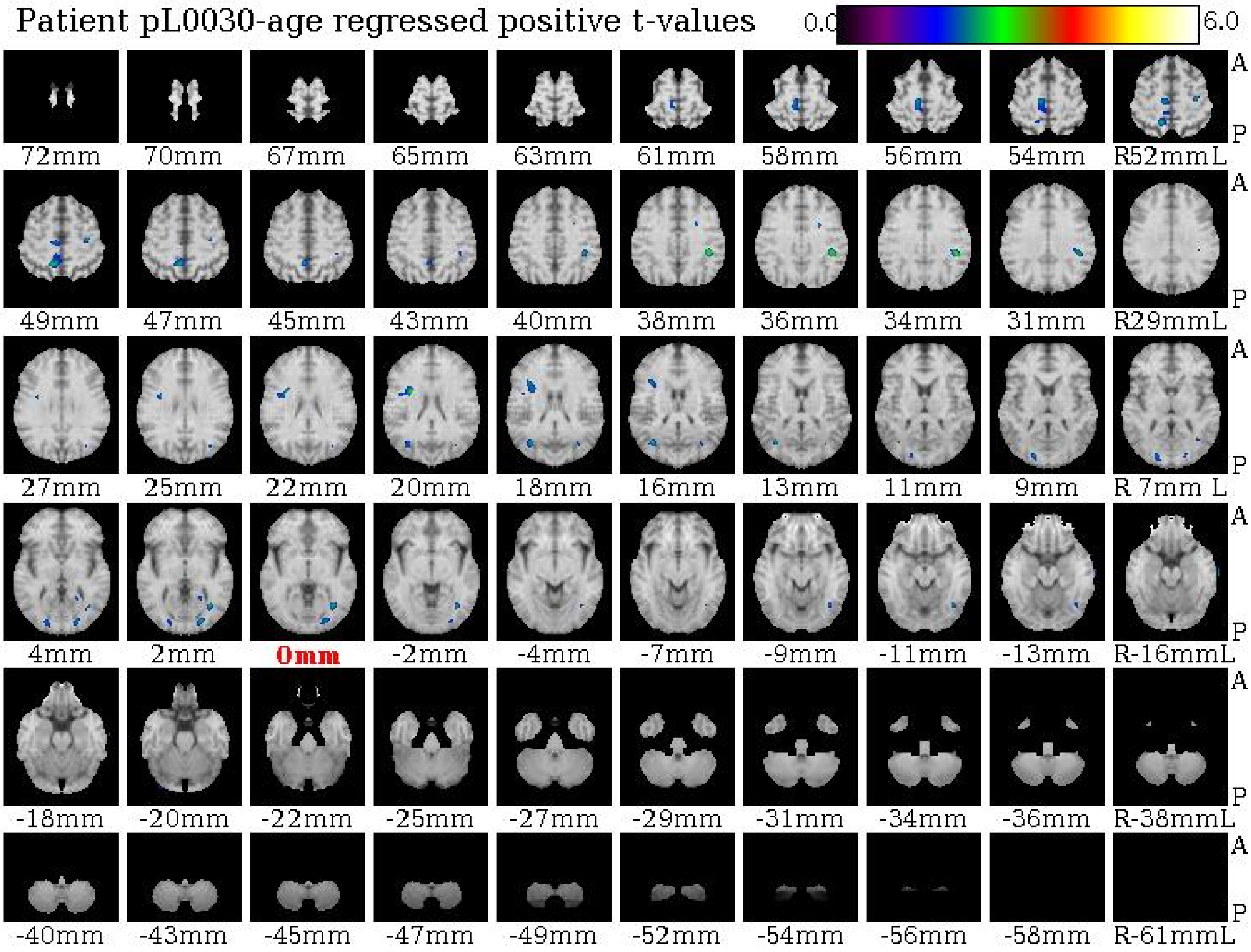

Patient pL0030-age regressed positive t-values
0.0
6.0
A
P
72mm
70mm
67mm
65mm
63mm
61mm
58mm
56mm
54mm
R52mmL
49mm
47mm
45mm
43mm
40mm
38mm
36mm
34mm
31mm
R29mmL
27mm
25mm
22mm
20mm
18mm
16mm
13mm
11mm
9mm
R 7mm L
4mm
2mm
0mm
-2mm
-4mm
-7mm
-9mm
-11mm
-13mm
R-16mmL
-18mm
-20mm
-22mm
-25mm
-27mm
-29mm
-31mm
-34mm
-36mm
R-38mmL
-40mm
-43mm
-45mm
-47mm
-49mm
-52mm
-54mm
-56mm
-58mm
R-61mmL

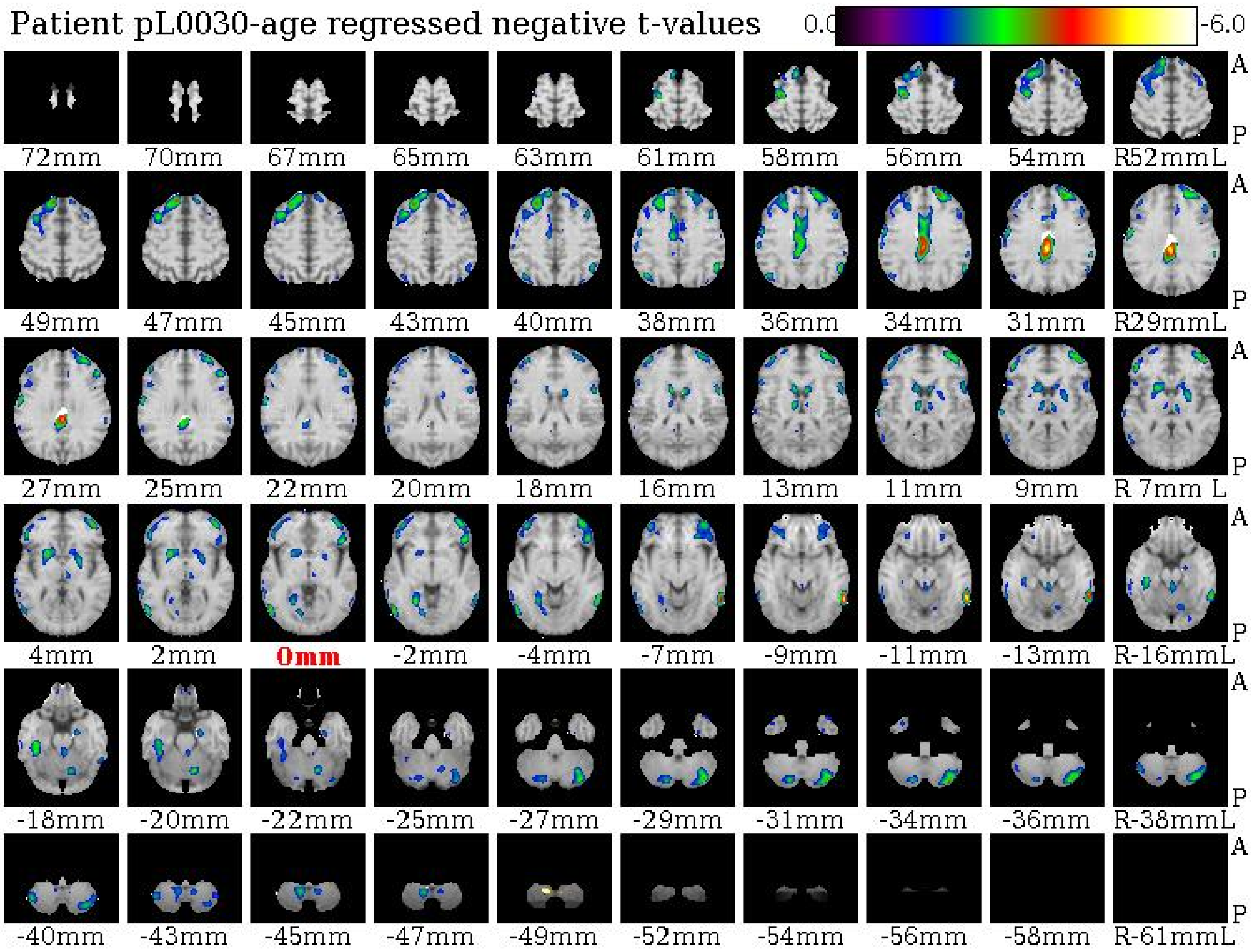

Patient pL0030-age regressed negative t-values
0.0
-6.0
A
P
72mm
70mm
67mm
65mm
63mm
61mm
58mm
56mm
54mm
R52mmL
49mm
47mm
45mm
43mm
40mm
38mm
36mm
34mm
31mm
R29mmL
27mm
25mm
22mm
20mm
18mm
16mm
13mm
11mm
9mm
R 7mm L
4mm
2mm
0mm
-2mm
-4mm
-7mm
-9mm
-11mm
-13mm
R-16mmL
-18mm
-20mm
-22mm
-25mm
-27mm
-29mm
-31mm
-34mm
-36mm
R-38mmL
-40mm
-43mm
-45mm
-47mm
-49mm
-52mm
-54mm
-56mm
-58mm
R-61mmL

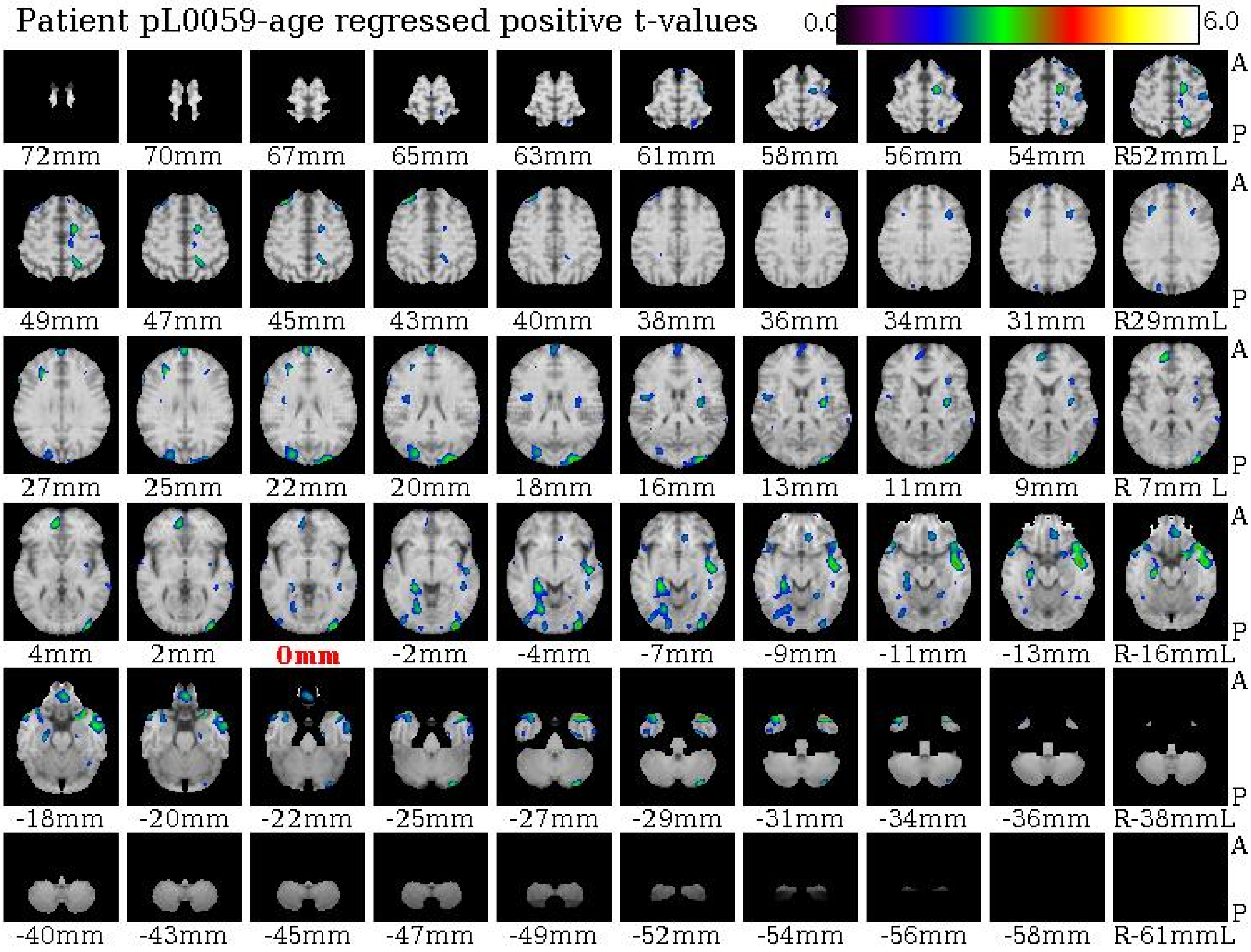

Patient pL0059-age regressed positive t-values
0.0
6.0
A
P
72mm 70mm 67mm 65mm 63mm 61mm 58mm 56mm 54mm R52mmL
49mm 47mm 45mm 43mm 40mm 38mm 36mm 34mm 31mm R29mmL
27mm 25mm 22mm 20mm 18mm 16mm 13mm 11mm 9mm R 7mm L
4mm 2mm 0mm -2mm -4mm -7mm -9mm -11mm -13mm R-16mmL
-18mm -20mm -22mm -25mm -27mm -29mm -31mm -34mm -36mm R-38mmL
-40mm -43mm -45mm -47mm -49mm -52mm -54mm -56mm -58mm R-61mmL

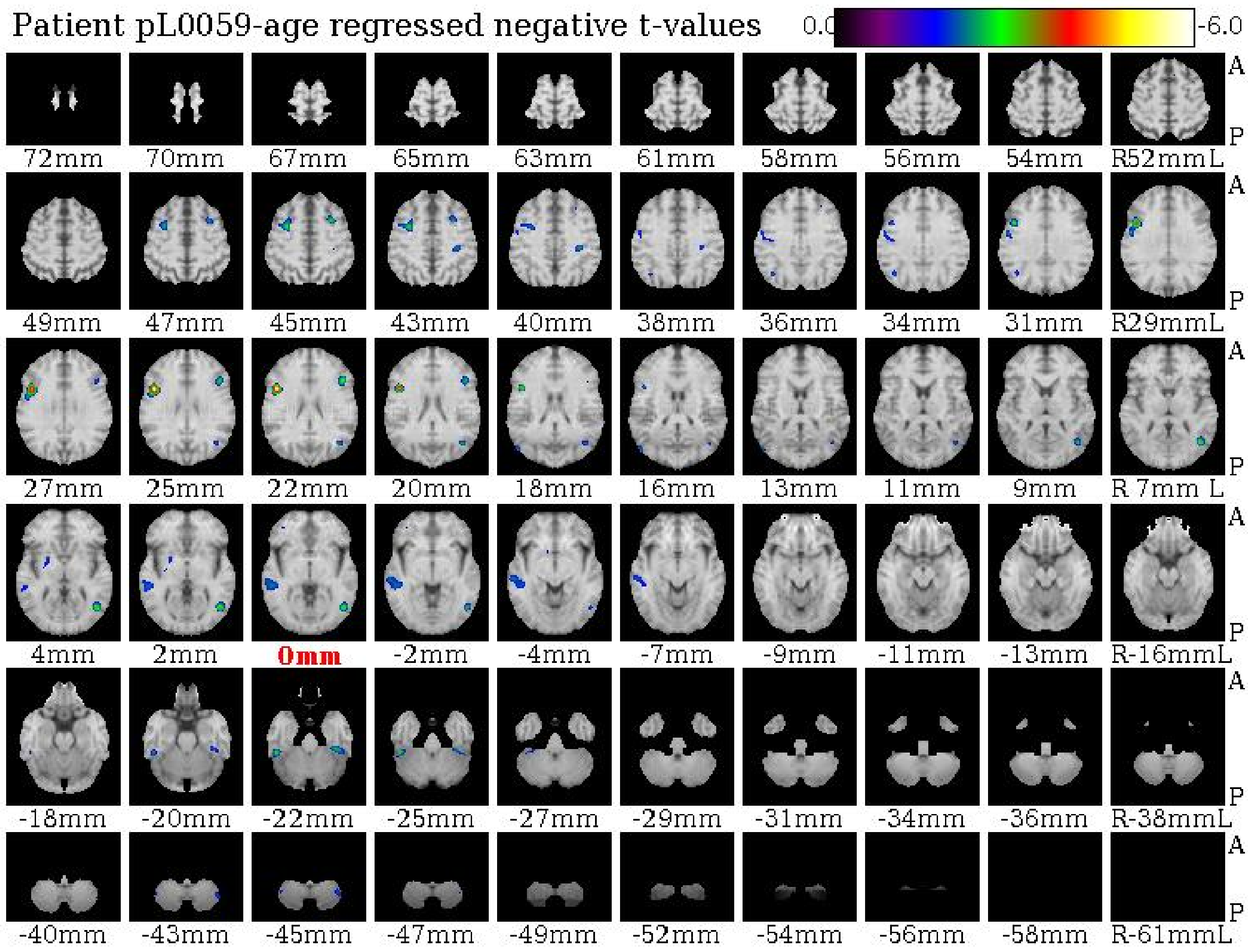

Patient pL0059-age regressed negative t-values
0.0
-6.0
A
P
72mm
70mm
67mm
65mm
63mm
61mm
58mm
56mm
54mm
R52mmL
49mm
47mm
45mm
43mm
40mm
38mm
36mm
34mm
31mm
R29mmL
27mm
25mm
22mm
20mm
18mm
16mm
13mm
11mm
9mm
R 7mm L
4mm
2mm
0mm
-2mm
-4mm
-7mm
-9mm
-11mm
-13mm
R-16mmL
-18mm
-20mm
-22mm
-25mm
-27mm
-29mm
-31mm
-34mm
-36mm
R-38mmL
-40mm
-43mm
-45mm
-47mm
-49mm
-52mm
-54mm
-56mm
-58mm
R-61mmL

Patient pL0071-age regressed positive t-values

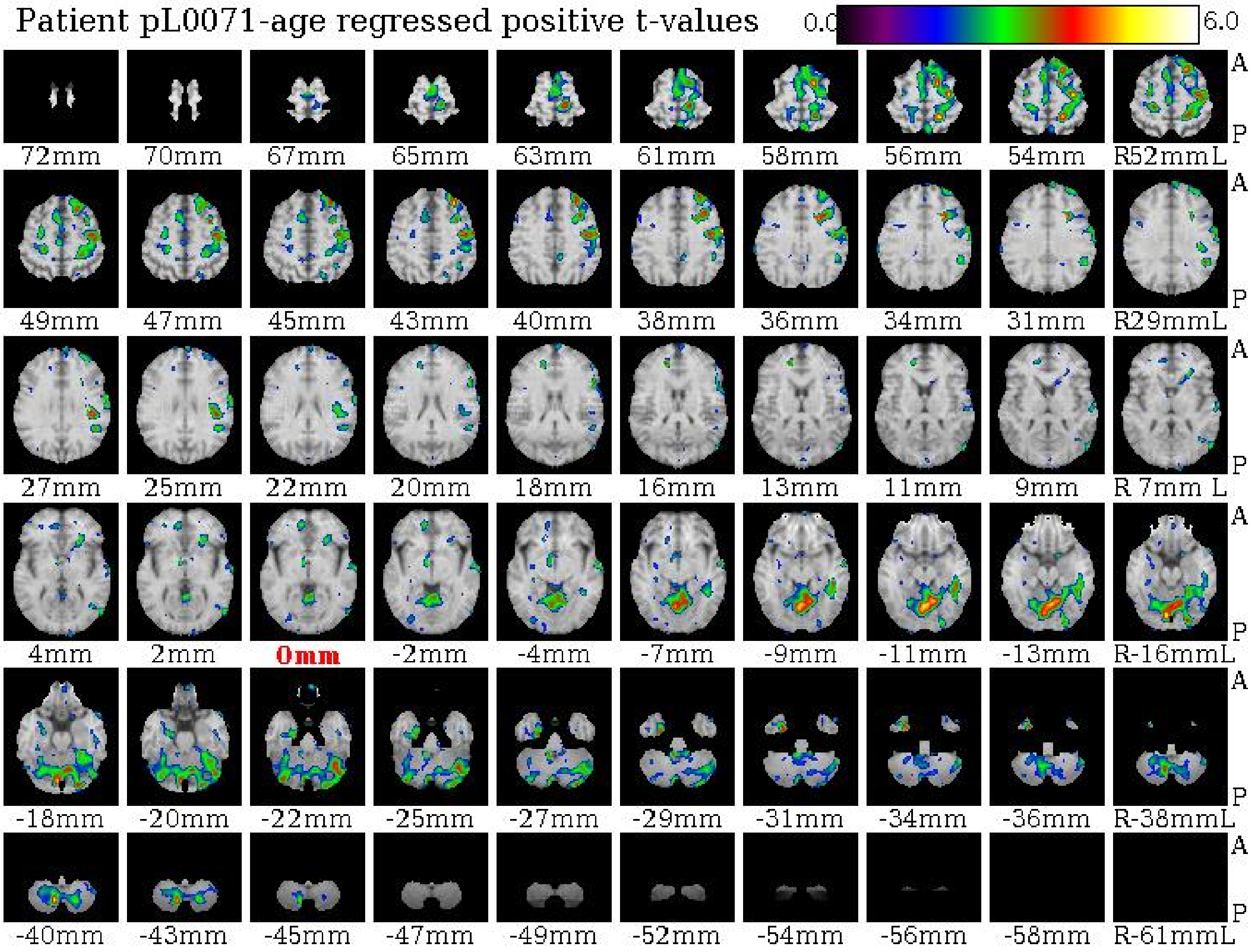

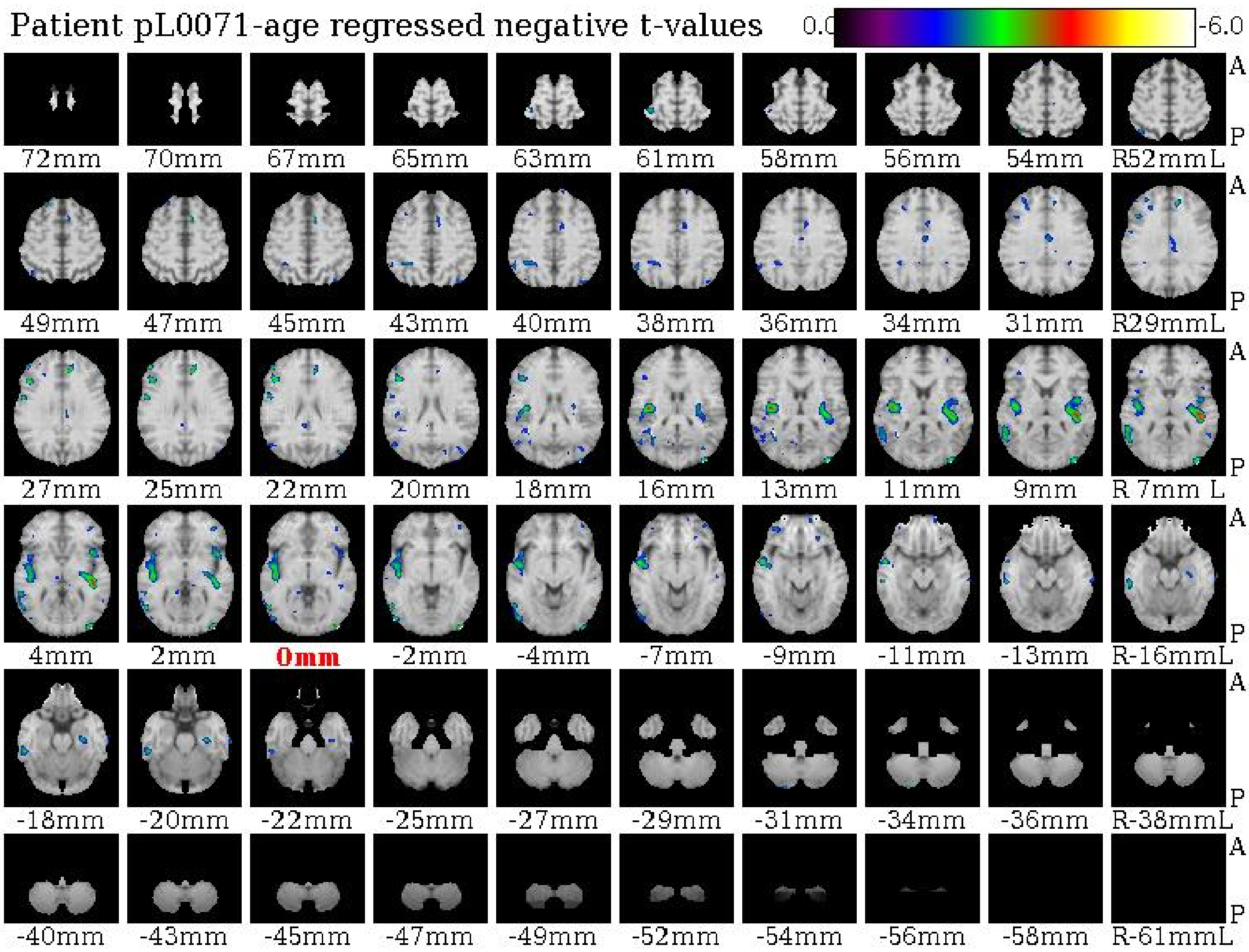

Patient pL0071-age regressed negative t-values
0.0
-6.0
72mm
70mm
67mm
65mm
63mm
61mm
58mm
56mm
54mm
R52mmL
49mm
47mm
45mm
43mm
40mm
38mm
36mm
34mm
31mm
R29mmL
27mm
25mm
22mm
20mm
18mm
16mm
13mm
11mm
9mm
R 7mm L
4mm
2mm
0mm
-2mm
-4mm
-7mm
-9mm
-11mm
-13mm
R-16mmL
-18mm
-20mm
-22mm
-25mm
-27mm
-29mm
-31mm
-34mm
-36mm
R-38mmL
-40mm
-43mm
-45mm
-47mm
-49mm
-52mm
-54mm
-56mm
-58mm
R-61mmL
A
P

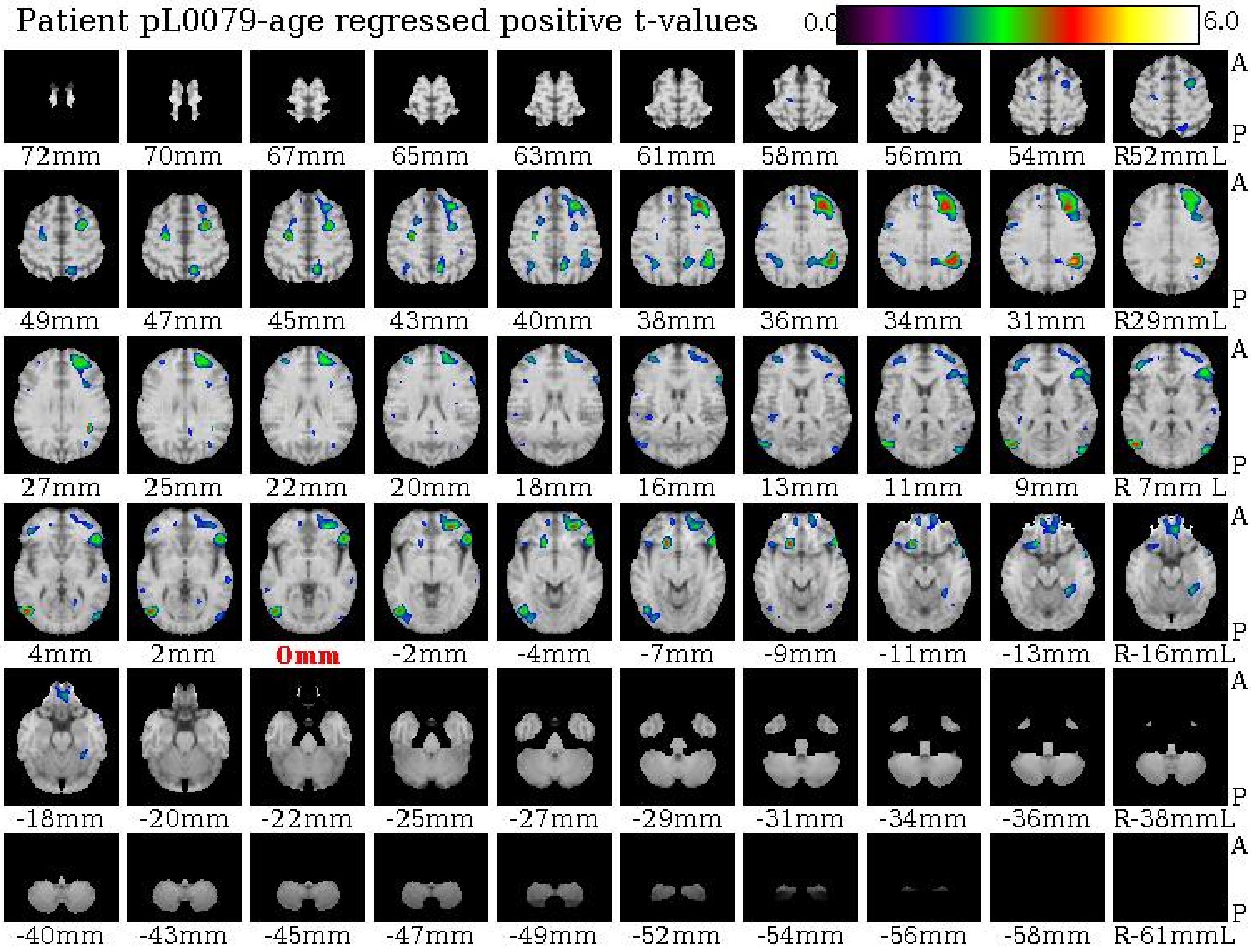

Patient pL0079-age regressed positive t-values
0.0
6.0
A
P
72mm
70mm
67mm
65mm
63mm
61mm
58mm
56mm
54mm
R52mmL
49mm
47mm
45mm
43mm
40mm
38mm
36mm
34mm
31mm
R29mmL
27mm
25mm
22mm
20mm
18mm
16mm
13mm
11mm
9mm
R 7mm L
4mm
2mm
0mm
-2mm
-4mm
-7mm
-9mm
-11mm
-13mm
R-16mmL
-18mm
-20mm
-22mm
-25mm
-27mm
-29mm
-31mm
-34mm
-36mm
R-38mmL
-40mm
-43mm
-45mm
-47mm
-49mm
-52mm
-54mm
-56mm
-58mm
R-61mmL

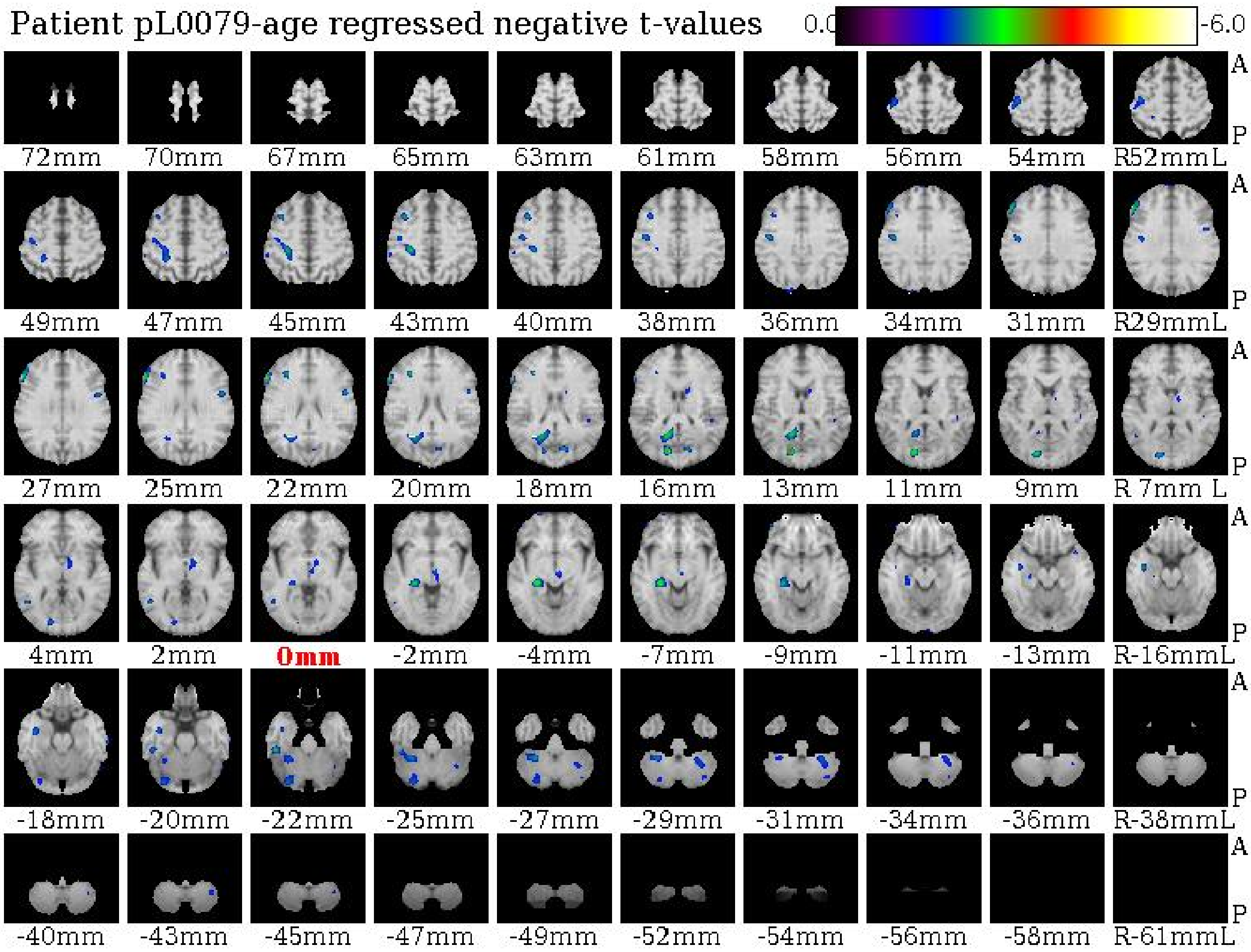

Patient pL0079-age regressed negative t-values
0.0
-6.0
A
P
72mm
70mm
67mm
65mm
63mm
61mm
58mm
56mm
54mm
R52mmL
49mm
47mm
45mm
43mm
40mm
38mm
36mm
34mm
31mm
R29mmL
27mm
25mm
22mm
20mm
18mm
16mm
13mm
11mm
9mm
R 7mm L
4mm
2mm
0mm
-2mm
-4mm
-7mm
-9mm
-11mm
-13mm
R-16mmL
-18mm
-20mm
-22mm
-25mm
-27mm
-29mm
-31mm
-34mm
-36mm
R-38mmL
-40mm
-43mm
-45mm
-47mm
-49mm
-52mm
-54mm
-56mm
-58mm
R-61mmL

Patient pL0089-age regressed positive t-values

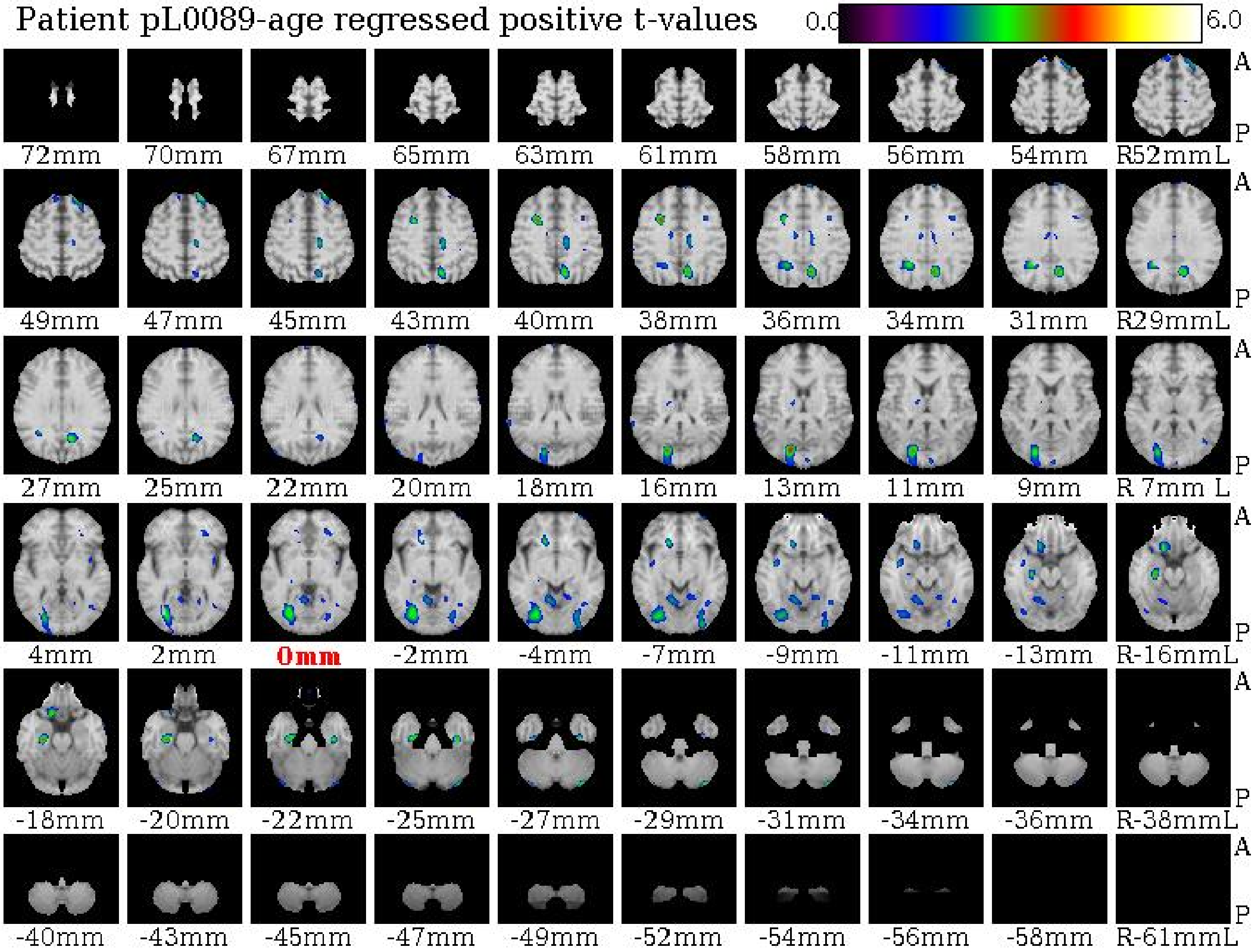

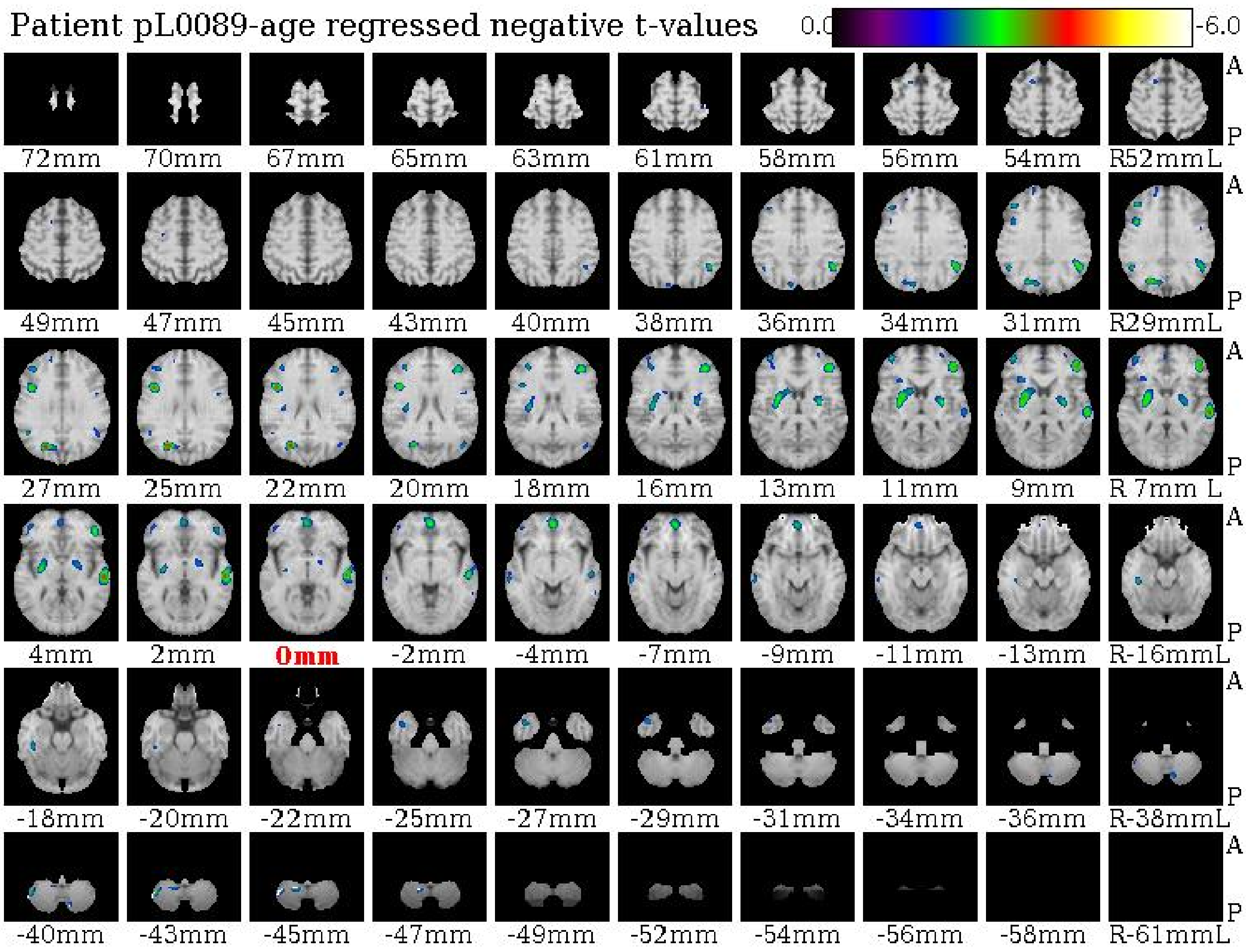

Patient pL0089-age regressed negative t-values
0.0
-6.0
A
P
72mm 70mm 67mm 65mm 63mm 61mm 58mm 56mm 54mm R52mmL
49mm 47mm 45mm 43mm 40mm 38mm 36mm 34mm 31mm R29mmL
27mm 25mm 22mm 20mm 18mm 16mm 13mm 11mm 9mm R 7mm L
4mm 2mm 0mm -2mm -4mm -7mm -9mm -11mm -13mm R-16mmL
-18mm -20mm -22mm -25mm -27mm -29mm -31mm -34mm -36mm R-38mmL
-40mm -43mm -45mm -47mm -49mm -52mm -54mm -56mm -58mm R-61mmL
